# Three-dimensional biofilm growth supports a mutualism involving matrix and nutrient sharing

**DOI:** 10.1101/2020.10.26.355560

**Authors:** Heidi A. Arjes, Lisa Willis, Haiwen Gui, Yangbo Xiao, Jason Peters, Carol Gross, Kerwyn Casey Huang

## Abstract

Life in a three-dimensional structure such as a biofilm is typical for many bacteria, yet little is known about how strains with different genotypes interact in this context. Here, to systematically explore gene knockdowns across various three-dimensional contexts, we created arrayed libraries of essential-gene CRISPRi knockdowns in the model biofilm-forming bacterium *Bacillus subtilis* and measured competitive fitness during colony co-culture with a wild type-like parent on different media and at different knockdown levels. Partial knockdown led to a wide range of fitness phenotypes, with targeting of translation-related genes often leading to lower growth rates and rapid out-competition by the parent. Several knockdowns competed differentially in biofilms versus non-biofilm colonies, in some cases due to lack of a particular nutrient in one medium. Cells depleted for the alanine racemase AlrA died in monoculture, but co-cultures survived via nutrient sharing in a biofilm but not in liquid. This rescue was enhanced in biofilm co-culture with a parent unable to produce extracellular matrix, due to a mutualism involving nutrient and matrix sharing. Including *alrA*, we identified several examples of mutualism involving matrix sharing that occurred in a three-dimensional biofilm colony but not when growth was constrained to two dimensions. These findings demonstrate that growth in a three-dimensional biofilm can promote genetic diversity through sharing of secreted factors, and illustrate the role of matrix production in determining trajectories for biofilm evolution that may be relevant to pathogens and other environmental bacteria.

## Introduction

In natural environments, many bacteria grow in dense, three-dimensional multicellular communities held together by extracellular matrix, often called biofilms. Biofilms have widespread clinical [1], industrial [2], and biotechnological [3] implications. Biofilms allow for genetic differentiation and division of labor that can mutually benefit distinct genotypes. For instance, in a dual-species biofilm, extracellular matrix components were functionally exploited by multiple species to drive emergent structural and mechanical properties of the biofilm that affected viability [4]. Additionally, the rate at which mutations fix in a population of a given size is higher in microbial colonies compared to well-shaken, liquid cultures [5], suggesting that spatial confinement supports an increase in genetic variation. Spatial confinement dramatically increases the frequency of interactions between nearby cells and thus the potential for coupled evolutionary outcomes, enhancing random genetic drift [6]. However, mechanisms that support genetic diversity in the context of a three-dimensional bacterial colony or biofilm remain underexplored [7].

The model organism *Bacillus subtilis* is a soil-dwelling species that adheres to plant roots as a biofilm [8]. In laboratory conditions, *B. subtilis* grows on surfaces in biofilm or non-biofilm colonies depending on the growth medium. In the commonly used rich medium LB, *B. subtilis* grows as a colony with limited biofilm characteristics [9]. By contrast, when cultured on the biofilm-promoting medium MSgg [10], *B. subtilis* natural isolates produce extracellular matrix composed of secreted polysaccharides and proteinaceous components that hold cells together and enhance biofilm colony expansion [7, 11, 12]. The extracellular matrix also determines colony architecture through the three-dimensional pattern of growth and wrinkling [13]. This architecture creates a variety of contexts for genetically identical cells to differentially express genes depending on their location, and indeed biofilms contain functionally distinct subpopulations [14, 15]: living cells differentiate into extracellular-matrix producers, sporulating cells, and motile cells, while dead cells may be cannibalized [16–18]. Thus, biofilms are an environment with heightened potential for interactions among cells in distinct transcriptional states and/or genetic backgrounds. Furthermore, biofilm-specific interactions can be identified and characterized by comparing biofilm and non-biofilm growth conditions.

The explosion of interest in microbial communities in recent years has stimulated a variety of approaches for identifying interspecies interactions. Liquid co-cultures have been used to quantify interaction networks [19] and to dissect changes in antibiotic sensitivity in co-cultures [20], but liquid growth cannot be used to identify biofilm or colony-specific interactions as it removes the spatial context of community growth and likely prioritizes long-range interactions over short-range physical contacts. Moreover, determining the amount of each strain in a co-culture often relies on laborious methods such as dilution plating and colony counting [20, 21], which may be complicated if cells adhere to each other. Microfluidics has facilitated the production and high-throughput analysis of droplets with mixed species, but these approaches again rely on liquid growth and the resulting community has very few cells compared to most natural communities [22, 23], making it difficult to study complex fitness phenotypes beyond those affecting initial growth. The production of antibiotics by certain species has been investigated using the inhibition of colony growth of other species at a distance [24–26], and a colony-based screen identified interspecies interactions between *B. subtilis* and other soil bacteria [27]. While powerful, these methods are not applicable to investigating genetically distinct strains growing together in a three-dimensional structure. The increased availability of mutant libraries across organisms [28–31] motivates the development of a colony-based strategy for high-throughput screening of the fitness of strains within co-cultures.

Both non-essential and essential genes (so defined based on survival in a typical laboratory environment such as liquid growth in LB) may impact fitness in any environment. While chemical-genetic screens of ordered libraries of deletions of non-essential genes have revealed novel phenotypes and elucidated the mechanism of action of drugs [32, 33], and phenotypic screens of transposon libraries have identified sporulation- and biofilm-related non-essential genes [34, 35], essential genes have traditionally been challenging to address. CRISPR interference (CRISPRi) utilizes an endonuclease-dead version of Cas9 (dCas9) to inhibit transcription from a gene of interest [36], facilitating tunable expression of any gene. Previously, we created a library of CRISPRi knockdowns of each essential gene in the non-biofilm-forming strain *B. subtilis* 168, which we used to uncover essential gene networks and to identify functional classes of genes based on growth and morphology [30]. In each strain of this library, the level of an essential gene can be titrated, from basal knockdown that allows robust growth of cells in liquid cultures, to full knockdown that inhibits the growth of many strains [30]. Thus, CRISPRi targeting of essential genes provides the potential for a wide distribution of phenotypes, enabling determination of the effects of essential gene disruption without completely inhibiting growth [37]. This ability to achieve tunable knockdown is particularly appealing for quantifying interstrain interactions, by contrast to the lethal phenotype of complete removal of essential genes. CRISPRi was recently used to identify genes that regulate biofilm formation in *Pseudomonas fluorescens* [38]; the efficacy of CRISPRi for *B. subtilis* in a colony/biofilm environment has yet to be ascertained.

Here, we created GFP-labeled libraries of CRISPRi essential-gene knockdowns in the biofilm-forming strain *B. subtilis* 3610 to investigate the fitness consequences of gene knockdowns and interstrain interactions within three-dimensional biofilm and non-biofilm colonies. We demonstrated that the level of CRISPRi knockdown is tunable during colony growth on LB and MSgg agar. We developed a high-throughput method for screening monocultures and co-culture colonies on agar plates, and applied this method to quantify growth and fitness when CRISPRi knockdowns were co-cultured with a wild-type-like parent strain. We observed a wide range of fitness phenotypes across media and knockdown levels, with partial knockdowns of translation-related genes producing the lowest fitness, likely due to their negative impact on growth rate. We discovered that full knockdown of *alrA*, which encodes an alanine racemase required for cell-wall synthesis, was rescued by the presence of wild-type cells in a co-culture biofilm colony but not in liquid. This rescue was enhanced and stable over time when parent cells were unable to produce extracellular matrix, revealing a mutualistic interaction between these strains. Finally, we identified several other knockdowns with higher competitive fitness when the parent cells are deficient in extracellular matrix production, as long as growth occurs in three dimensions, suggesting that these genes have mutualistic potential via nutrient and matrix sharing. These findings highlight the importance of colony geometry and matrix production in determining gene essentiality and interstrain genetic interactions, and provide foundational knowledge of mechanisms that support genetic diversity in in pathogenic and environmental biofilms.

## Results

### Construction of a knockdown library for probing gene essentiality in *B. subtilis* 3610

To study genetic interactions involving critical cellular processes within a biofilm, we constructed a CRISPRi knockdown library in the biofilm-forming *B. subtilis* strain 3610 (Methods). The library contains 302 strains: the 252 known essential genes in *B. subtilis* strain 168, 47 genes that were initially classified as essential in 168 [39] but later revealed to be non-essential or conditionally essential [29], and three internal controls expressing dCas9 without any guide RNAs (Table S1) [30]. Each strain in the library contains a xylose-inducible copy of *dcas9* and an sgRNA targeting the gene of interest (Fig. 1A). In addition, *gfp* is incorporated at the *sacA* locus to allow visualization and quantification of the knockdown strain (Fig. 1A). The *sacA::gfp* strain exhibited similar growth and biofilm wrinkling as a parental unlabeled control on both non-biofilm LB agar and biofilm-promoting MSgg [10] agar (Fig. 1B). We refer to colonies on LB and MSgg as “non-biofilm” colonies and “biofilm” colonies, respectively, although it is important to note that the biofilm definition is nuanced and colonies on LB may have some biofilm characteristics [9].

**Figure 1:**
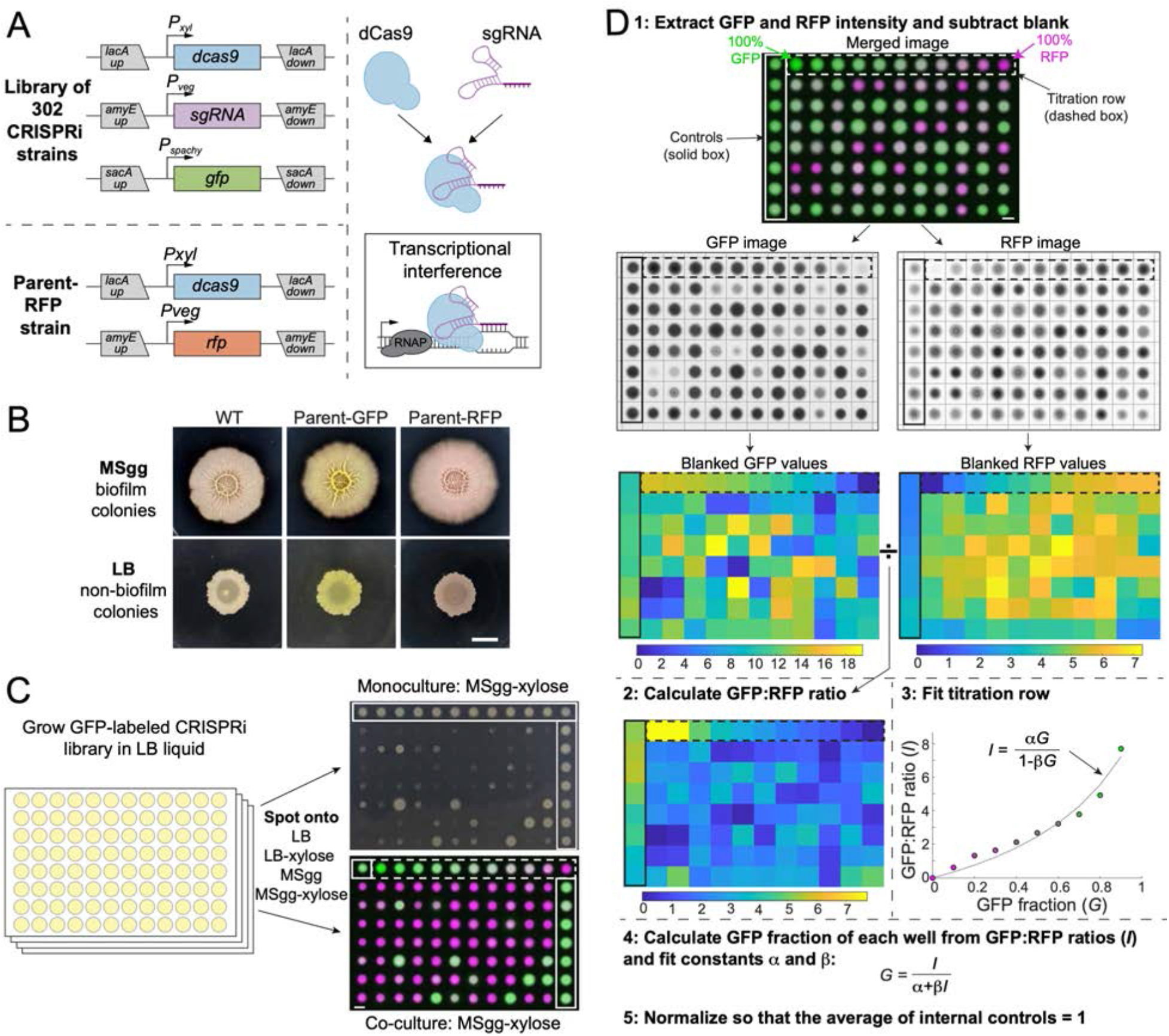
A high-throughput screening strategy to measure colony-based competition within bacterial colonies. A) We constructed a GFP-labeled library of CRISPRi knockdowns of all known essential and conditionally essential genes (top left). In the library, the nuclease-deactivated Cas9 gene (*dcas9*) is inducible with xylose and the single-guide RNA (*sgRNA*) is constitutively expressed. dCas9 binds the sgRNA and blocks transcription by physically impeding RNA polymerase (right). Every strain is labelled with *gfp* expressed from the *sacA* locus. A parent strain (parent-RFP, bottom left) that expresses *rfp* as well as *dCas9* without an *sgRNA* was used in competition assays. B) The parent-GFP (*sacA::gfp, lacA::dCas9*) and the parent-RFP strains have similar phenotypes to wild type on both biofilm-promoting MSgg agar and non-biofilm-promoting LB agar. Cultures were grown in liquid LB to an OD_600_~1 and then 1 μL was spotted in the middle of wells of a 6-well plate containing LB agar or MSgg agar. Image intensities were adjusted identically; the yellow and red colors of the parent strains are due to GFP and RFP expression, respectively. Scale bar: 5 mm. C) Schematic of screening strategy to measure the monoculture colony size and competitive fitness of each knockdown against the parent-RFP strain. GFP-labeled knockdown libraries were grown in liquid culture in 96-well microtiter plates. Monocultures were spotted onto LB and MSgg agar plates (top right) without or with xylose to achieve basal or full knockdown, respectively, of the targeted gene. The monocultures contain parent controls in wells along an outer column and row of the plate (solid box). Co-cultures of a 1:1 volumetric mixture of the parent-RFP and GFP-labeled library strains were spotted onto agar plates of LB and MSgg, without or with xylose. Controls in which parent-GFP was mixed with parent-RFP are bounded by horizontal red box. Bottom right: merged image of RFP and GFP signals from co-cultures. The co-cultures include a titration row from 100% GFP cultures to 100% RFP cultures in 10% increments (dashed box), and several controls of 1:1 mixtures of the parent GFP and parent-RFP strains (purple box). Scale bar: 5 mm. D) Schematic of image analysis to quantify competitive fitnesses from the co-culture screen. Data from plate 1 spotted on MSgg is presented as an example. Plates were segmented and individual colony intensities were extracted from the GFP and RFP images. GFP intensities were divided by RFP intensities to obtain ratios *I*. The titration row (dashed box) was fit to a curve using the equation *I*=α*G*/(1-β*G*), where *G* is the fraction of the parent-GFP strain, to extract fit parameters a and β for each plate individually. These parameters were used to map the GFP fractions of each colony and values were normalized so that the parent-GFP:parent-RFP control co-cultures on each plate (solid box) had an average value of 1. Scale bar: 5 mm.

To determine whether CRISPRi can be used to knock down gene expression in non-biofilm and biofilm colonies, we engineered a parent strain containing *rfp* under a constitutive promoter and used CRISPRi to target *rfp*. In this strain, RFP levels in non-biofilm colonies on LB agar plates ranged from 40% (basal knockdown) to ~0% (full knockdown) (Fig. S1A), a comparable range to knockdown of the domesticated strain 168 in liquid LB [30]. RFP levels in biofilm colonies on MSgg agar ranged from ~90% (basal knockdown) to ~0% (full knockdown) (Fig. S1A). Thus, CRISPRi is an effective tool to knock down gene expression in non-biofilm and biofilm colonies.

### High-throughput screening of competitive fitness in a colony

We compared the colony-growth phenotypes of GFP-labeled knockdown strains grown either alone or mixed with a control strain modified with xylose-inducible dCas9 (without an sgRNA) and constitutive expression of RFP (henceforth referred to as parent-RFP) that exhibits wild-type-like biofilm formation (Fig. 1A,B). After growing each strain individually in liquid LB, GFP-labeled knockdown strains were spotted either alone or mixed with parent-RFP onto agar plates (Fig. 1C, Fig. S2). Colony phenotypes were quantified using a custom image-analysis pipeline that segmented plates into colonies and computed the ratio of GFP:RFP for each colony; colony size was measured manually (Fig. 1D, S1B,C; Methods). Each plate included a titration row of colonies grown from mixtures of the parent-GFP strain with the parent-RFP strain at known concentrations from 0% to 100% parent-GFP (Fig. 1D, Fig. S2). Quantification of the titration row closely agreed with the predicted ratio of GFP:RFP at each time point (16, 24, and 48 h) (Fig. 1D, S1D), indicating that the relative fraction of GFP-labeled mutants in co-culture with the parent-RFP strain can be accurately quantified through comparison of the GFP:RFP ratio with the titration row (Methods). Thus, our screen allows us to quantitatively compare growth as a monoculture to growth in co-culture through this competitive fitness value (Fig. 1C).

### Gene knockdown results in a broad range of colony sizes and competitive fitnesses

To measure growth in monocultures or co-cultures across conditions, we spotted knockdowns on their own or mixed with parent-RFP on LB and MSgg agar without and with xylose. After 16 h of growth, colony sizes of basal knockdown monocultures exhibited a narrow distribution on LB agar, but were more widely distributed on MSgg agar (Fig. 2A, Fig. S3A, Table S2). By contrast to growth as monocultures on LB, basal knockdowns co-cultured with the parent for 16 h showed a broadly distributed competitive fitness on LB agar: only 92 of the strains had a fitness within 2 standard deviations of the mean of controls, while the remaining 210 strains were significantly defective in competition (below 2 standard deviations of the mean of the controls) (Fig. 2B, Table S2). A competitive fitness of 1 signifies equal amounts of GFP-labeled knockdown and parent-RFP, and 0 means that the GFP-labeled knockdown was completely out-competed by the parent-RFP strain. On MSgg agar, competitive fitness displayed a similar trend, with 198 basal knockdowns exhibiting a significant fitness defect after 16 h (Fig. 2A,B, Table S2). As expected, when transcription was fully knocked down, fitness was even more compromised: 168 and 143 strains had fitness <0.08 after 16 h of growth on LB and MSgg agar, respectively (Fig. 2B, Table S2). Together, these data demonstrate that even though phenotypes were generally subtle for monocultures grown on non-biofilm-promoting LB agar, screening the library on biofilm-promoting MSgg agar or in competition with a parent strain uncovered phenotypes even under basal knockdown.

**Figure 2:**
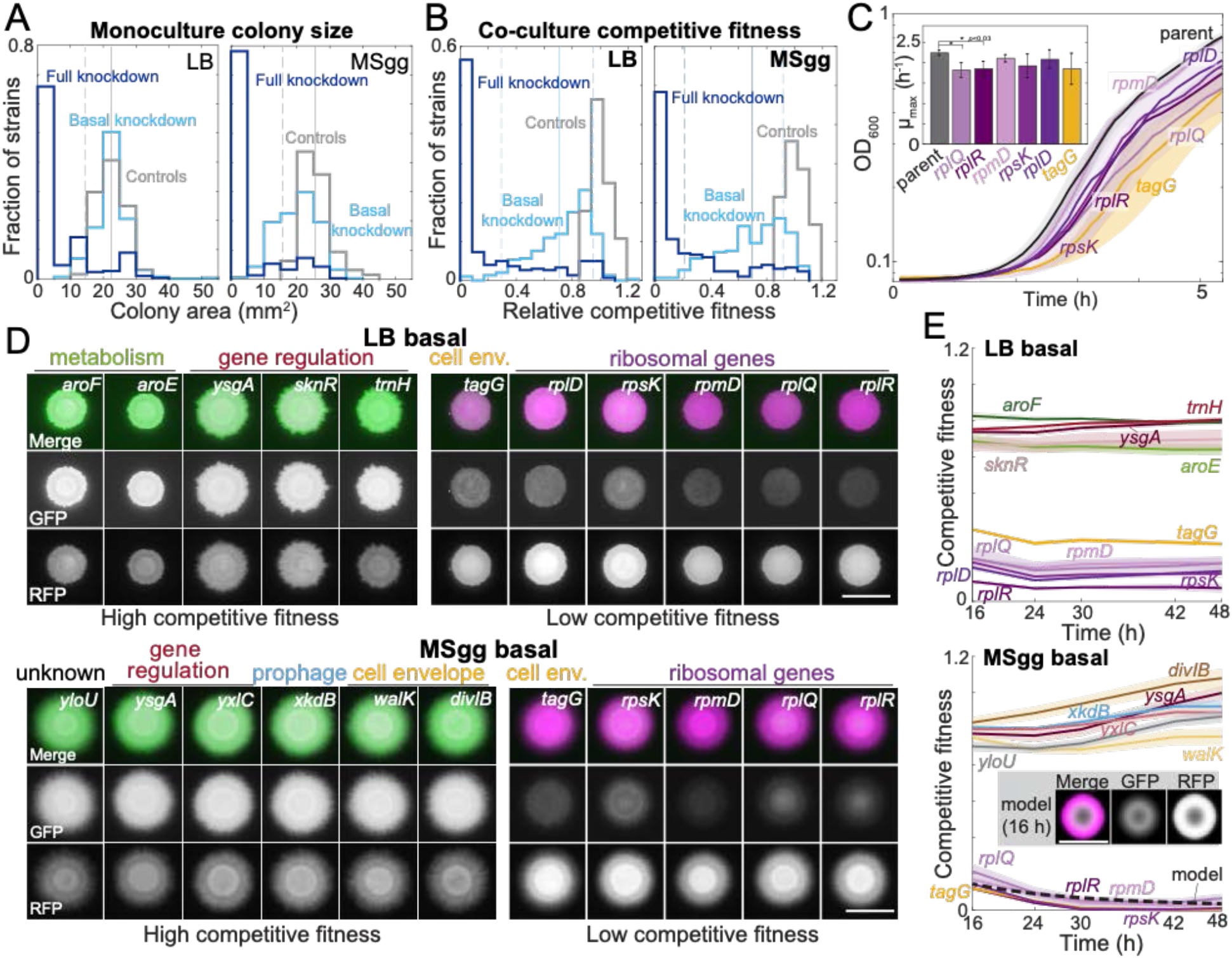
Growth on biofilm-promoting medium, increased knockdown, and competition against parent-RFP all broaden the distribution of fitnesses across the library. A) Basal knockdown (light blue) of essential genes on LB agar (which does not promote biofilms) resulted in similar colony sizes as parent-GFP controls (gray); only 13 of 302 colonies had size two standard deviations below the mean of the controls. By contrast, on biofilm-promoting MSgg agar the distribution of colony sizes spread to smaller values, with 80 colonies more than 2 standard deviations below the mean of the controls. Full knockdown (dark blue) inhibited growth of most strains. Data are from measurements at 16 h using the *sacA::gfp* library. Vertical solid lines show the mean of the control distribution and dashed lines show two standard deviations below the mean. B) 17 (LB) and 11 (MSgg) knockdown strains in the library competed poorly against the parent-RFP strain at basal knockdown (light blue), while 41 (LB) and 46 (MSgg) had competitive fitness similar to parent-GFP+parent-RFP controls (gray) even at full knockdown (dark blue). Data are from competition ratios at 16 h using the *sacA::gfp* library. Low-fitness strains were defined as having fitness two standard deviations below the mean of the data, and neutral-fitness strains were defined as having fitness above one standard deviation of the mean of the data. Vertical solid lines show the mean and dashed lines show two standard deviations below the mean and one standard deviation above the mean of the data from basal knockdown. C) Strains with low competitive fitness for basal knockdown generally had lower growth rate in liquid monoculture than parent control strains. Colonies were inoculated into liquid LB and OD_600_ was monitored over time. Ribosome-related genes are shown in shades of purple and a cell envelope-related gene (*tagG*) is shown in yellow. Curves are means and shaded regions represent 1 standard deviation (*n*=3). Inset: maximum growth rates. *: *p*<0.03, Student’s unpaired *t*-test, Benjamini-Hochberg multiple test correction applied. D) On both LB and MSgg agar, basal knockdown of *ygsA*, which is involved in gene regulation, exhibited high competitive fitness (left) and *tagG* and ribosomal-gene knockdowns exhibited low competitive fitness (right). GFP (knockdown strain) is false-colored green and RFP (parent) is false-colored magenta. Images are from 16 h using the *sacA::gfp* library. Scale bar: 5 mm. E) Competitive fitness of the strains with the highest and lowest values was approximately constant after 16 h. Curves are means and shaded regions represent the standard error of the mean (*n*=3 independent measurements). Inset and dashed black line: A reaction-diffusion model of co-culture colony growth with physically realistic parameters indicates that knockdowns (magenta) with maximum growth rate 20% lower than the parent (green) reproduces the colony sizes (bottom, inset) and competitive fitness (bottom, dashed black line) of ribosomal protein knockdowns after 16 h (Methods, Fig. S3C).

Several strains competed poorly with the parent even with basal knockdown in both non-biofilm and biofilm colonies (Fig. 2B). Interestingly, some non-essential genes had low competitive fitness. For instance, *mapA*, which encodes a methionine aminopeptidase, competed poorly in both LB and MSgg colonies, and CRISPRi induction further reduced fitness (Table S2, Fig. S3B). Analysis of DAVID functional annotations of strains with competitive fitness >2 standard deviations below the mean of controls revealed significant enrichment of structural constituents of ribosomes (*p*=9.8×10^−4^ and *p*=2.1×10^−2^ on LB and MSgg agar, respectively). Some of the ribosomal-protein strains that competed most poorly exhibited ~20% lower maximum growth rate than wild type in liquid cultures (Fig. 2C), suggesting that the reduced competitive fitness of these strains is due to their reduced growth rate. Indeed, a reaction-diffusion model of colony growth of a co-culture indicated that a strain’s maximum growth rate is a major determinant of competitive fitness, and that the 20% decrease of maximum growth rate in certain ribosomal protein knockdowns is consistent with our experimental measurements of their competitive fitness (Fig. S3C, Methods).

By contrast, several strains had fitness in co-culture with the wild-type-like parent similar to controls for basal and/or full induction (Fig. 2B), suggesting that the targeted gene was rendered less essential by co-culture with the wild-type-like parent. DAVID functional enrichment analysis of basal knockdowns with fitness values within 1 standard deviation of the mean across controls (*n*=41 and 46 strains for LB and MSgg, respectively) highlighted integral membrane components on both solid media (*p*=2.1×10^−3^ for both LB and MSgg). Under basal conditions, there were 4 knockdowns (*menH* and *cytC* on LB and *aroF* and *rny* on MSgg) in which monoculture growth was clearly compromised by induction but competitive fitness remained high (Fig. S3B). As *menH* and *aroF* are involved in synthesis of menaquinone (vitamin K2) and aromatic amino acids, respectively, the high competitive fitness may result from nutrient sharing within the colony. In addition, aromatic amino acid biosynthesis genes were enriched in basal knockdowns with high competitive fitness on LB agar (*p*=1.6×10^−2^), potentially due to the presence of aromatic amino acids in rich LB medium but not in MSgg. These data underscore the medium-dependence of gene essentiality in co-culture colonies.

To validate these findings, we replicated fitness measurements over time on a subset of strains with the highest or lowest competitive fitness values during basal knockdown on LB or MSgg agar. We found that the competitive fitness phenotype was highly reproducible and relatively stable over two days of colony growth (Fig. 2D,E, S3D,E), highlighting the utility of our CRISPRi library for probing the fitness of essential gene knock down in co-cultures.

### Several biosynthesis-related genes have different phenotypes in rich versus minimal media

The distinct nutrient compositions of LB and MSgg, along with the much broader distribution of monoculture colony sizes in MSgg compared with LB (Fig. 2A), motivated a comparison of competitive fitness across media. Somewhat surprisingly given the likelihood of different metabolic profiles due to media compositional differences, 93% of the strains exhibited similar competitive fitness (defined here as within 0.24 or 0.3 of the *y=x* line for basal or full knockdown, respectively) on MSgg and LB agar, whether the targeted gene was basally or fully knocked down (Fig. 3A, Table S2). Nonetheless, we identified 36 strains competed with the parent-RFP strain better on MSgg than on LB, and/or vice versa, under basal or full knockdown (Fig. 3A,B, Table S3). Strains that competed better in one medium compared to the other were statistically enriched for genes involved in amino-acid biosynthesis (*p*=4.7×10^−6^ and *p*=8.3×10^−5^ for basal and full knockdown, respectively), suggesting that some strains benefit from nutritional components specific to one medium (Fig. 3A, Table S3).

**Figure 3:**
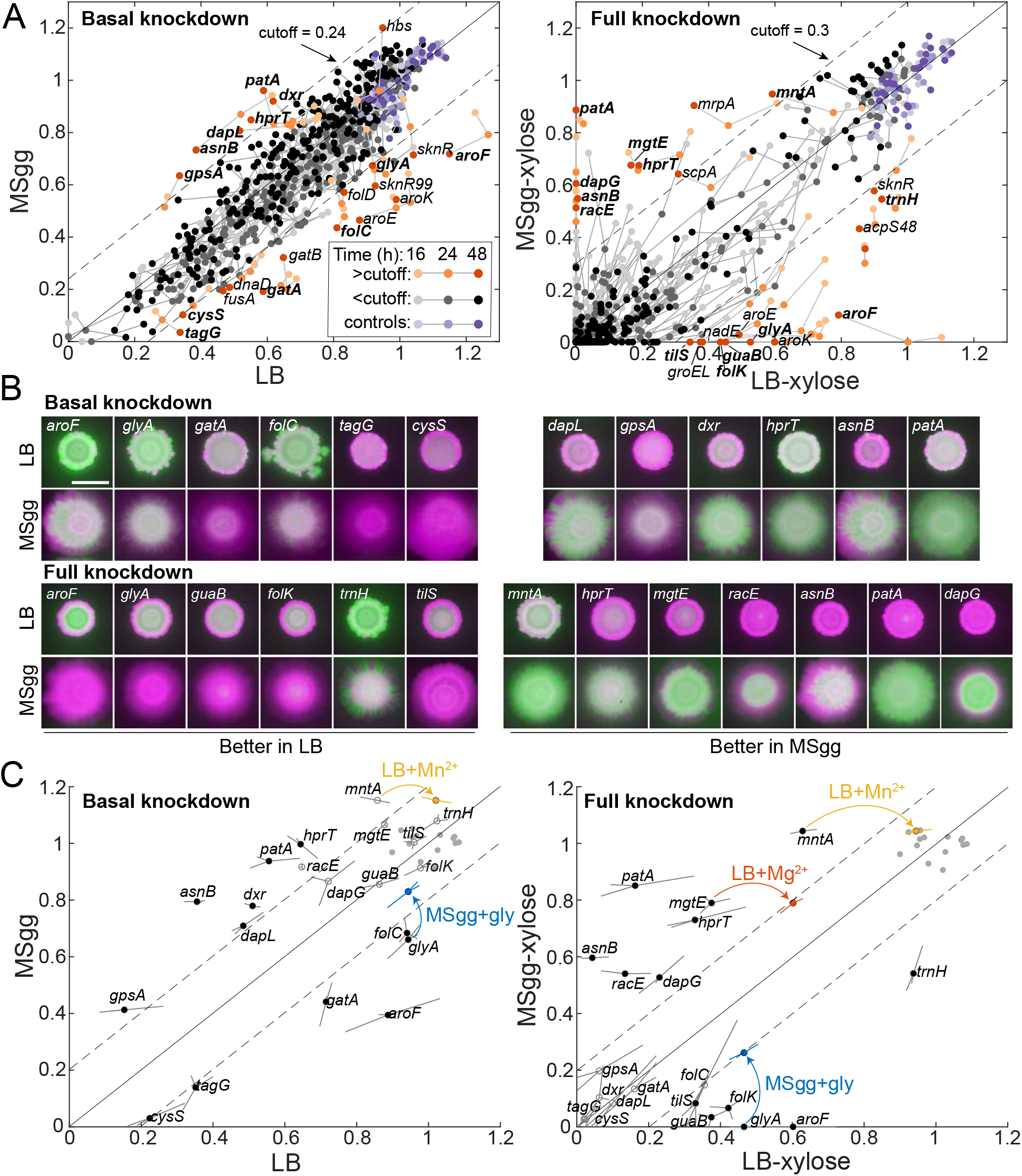
Some media-specific differences in competetive fitness can be directly attributed to the nutrient composition of the media. A) Although most knockdowns had similar fitness on LB and MSgg agar, a subset of knockdowns had higher competitive fitness on LB agar than on MSgg agar, or vice versa. Genes with fitness difference >0.24 at 24 h or >0.3 at 48 h are annotated and colored in orange shades, while those below the cutoff are in grayscale and parent-RFP+parent-GFP co-culture controls are in shades of purple. Genes labeled in bold were selected for follow-up studies. Data are from the *sacA::gfp* library at 16, 24, and 48 h. The solid line is *y=x* and the dotted lines represent the chosen cutoff. B) Images of colonies of the bolded genes in (A) after 48 h that illustrate the differential competitive fitness between LB and MSgg. Green and magenta represent fluorescence from the gene knockdown and parent, respectively. Scale bar: 5 mm. C) Addition of specific nutrients to the medium with poorer competitive fitness rescued competitive fitness for the *mntA, glyA*, and *mgtE* knockdowns. Means (filled circles) of triplicates (shown at end of lines extending from the circle) are displayed. Parent-RFP+parent-GFP controls are shown as gray filled circles. Data from addition experiments (LB+manganese, LB+magnesium, and MSgg+glycine for *mntA, mgtE*, and *glyA*, respectively) are shown as colored circles and lines at the ends of arrows. All changes marked with arrows are significant after correcting for multiple hypotheses with the Benjamini-Hochberg method (*p*<0.01, Student’s unpaired *t*-test).

Despite the undefined nature of LB, it was still possible for many strains to identify candidate components whose addition to the medium with lower competitive fitness might rescue the deficit. We selected 20 knockdowns with medium-dependent fitness to pursue further, many of which displayed fitness differences for both basal and full knockdown (Table S3). The *glyA* knockdown had higher competitive fitness on LB agar than on MSgg agar, and as hypothesized, addition of glycine to MSgg significantly improved fitness in basal and induced conditions, to levels closely matching fitness on LB (Fig. 3C). Similarly, adding Mg^2+^ or Mn^2+^ to LB agar at the same levels as in MSgg restored the competitive fitness of *mgtE* and *mntA* full knockdowns, respectively, to levels on MSgg (Fig. 3C, S4A). Somewhat surprisingly, even though many of the remaining 17 knockdowns with medium-dependent fitness naturally suggested candidates for a missing nutrient, they did not exhibit increased fitness when the hypothesized nutrient was added to the medium with reduced fitness (Fig. S4B, Table S3), showing that at least exogenous provision of those nutrients is insufficient to complement the medium-specific fitness defect. Together, these results indicate that medium-dependent competition ratios can arise due to both nutrient compositional differences between media and other mechanisms that remain unindentified but highlight potentially important factors in selection during colony growth.

### Wild-type cells rescue *alrA* knockdown in a biofilm colony by sharing D-alanine

Since growth in a structured community provides opportunities for nutrient sharing and cellular differentiation, we hypothesized that some essential-gene knockdowns would be unable to grow as a colony in monoculture but would fare better in co-culture with wild-type-like parent-RFP cells. Across our *sacA::gfp* essential-gene knockdown library, we did not identify any knockdowns that exhibited robust growth in a colony co-culture but died as a monoculture (Fig. S2, S3, Table S2). (To conform to our inoculation protocol for biofilm cultures in which we used 1 μL of a liquid culture of OD ~1.0 (Methods), our high-throughput screen involved inoculation of each ~2 mm-diameter spot with ~2×10^5^ cells, likely facilitating the partial growth of some knockdowns that would be hampered in growth from a single cell on plates with xylose.) We found that disruption of the *thrC* locus prevents wrinkling formation on MSgg, presumably reflecting a growth defect, so we hypothesized that insertion of *gfp* at the *thrC* locus might exacerbate growth inhibition due to knockdown of certain essential genes. Thus, we constructed a second, *thrC::gfp* library of knockdowns of all 302 strains and screened it on MSgg with xylose. In the *thrC::gfp* library, *alrA* was the only knockdown that failed to form a colony as a monoculture but survived with the parent-RFP strain in biofilms on MSgg agar with xylose (Fig. 4A, S5A,B). AlrA is a racemase that converts L-alanine to D-alanine and is required for cross-linking of the peptidoglycan cell wall (Fig. 4B). Full knockdown of *alrA* expression during liquid growth in a strain without *gfp* disruption of *thrC* led to bulging indicative of cell-wall defects (Fig. 4C, S5C). The *alrA* strain from the *thrC::gfp* library managed to grow as a monoculture into a colony similar in size to the inoculation spot on LB-xylose agar but not beyond, as did the *alrA* strain from the *sacA::gfp* library on LB-xylose and MSgg-xylose plates. As discussed above, the absence of complete lysis is likely due to the high initial density driving growth of a visible colony (Fig. S3A); streaking all *alrA* strains resulted in a substantial reduction in the number of colonies on plates with xylose compared to without for both LB and MSgg (Fig. S5D), indicating that the full-knockdown phenotype across all media and genotypes is severe for *alrA* at lower initial cell densities.

**Figure 4:**
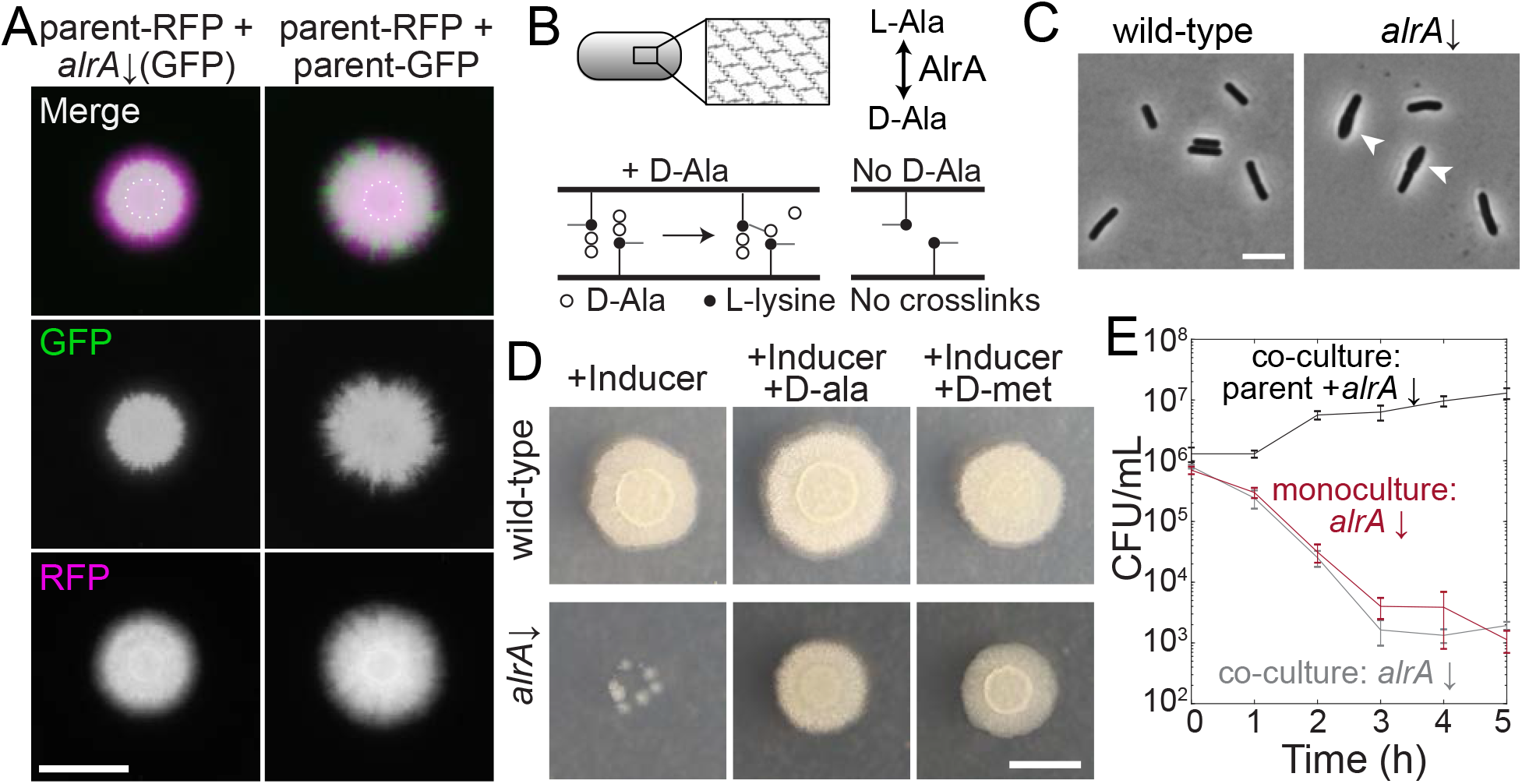
Full knockdown of *alrA* is rescued in a biofilm by D-alanine nutrient sharing, but not in liquid culture. A) Left: the *sacA::gfp alrA* knockdown under full induction was rescued by growth with the parent-RFP strain under biofilm-promoting conditions (MSgg agar). The *alrA* knockdown expanded beyond the boundaries of the original inoculum (dashed circle) when grown in co-culture with the parent-RFP strain. Right: the control co-culture of parent-RFP with parent-GFP preserves both strains at approximately equal proportions. Images were acquired at 24 h. In merged images, GFP from the *alrA* knockdown is false-colored green and RFP from the parent-RFP strain is false-colored magenta. Scale bar: 5 mm. B) AlrA is a racemase that converts L-alanine to D-alanine. D-alanine is critical for cell-wall crosslinking. C) Full knockdown of *alrA* caused cells to bulge, signifying cell-wall defects. Cells were cultured for 6 h in liquid MSgg with xylose to fully inhibit *alrA* expression. Arrowheads indicate bulging cells. Scale bar: 5 μm. D) Full knockdown of *alrA* was rescued by exogenous D-alanine. Cultures were grown in liquid LB to an OD_600_~1 and then 1 μL was spotted on MSgg agar alone or supplemented with 0.04 mg/mL D-alanine or D-methionine. Cells from *alrA* monocultures mostly died (left); the small colonies represent suppressors present in the initial inoculum. By contrast, addition of D-alanine (middle) or D-methionine (right) resulted in comparable growth to wild-type. Images are of an unlabeled *alrA* knockdown (HA420) and were acquired after 24 h of growth. Scale bar: 5 mm. E) Full knockdown of *alrA* was not rescued when co-cultured with the parent-RFP strain in liquid. For the co-culture, parent and *alrA* knockdown cultures were mixed 1:1 and back-diluted 1:100 into liquid MSgg with xylose to fully deplete *alrA*. For the *alrA* knockdown monoculture, the culture was diluted 1:200 into liquid MSgg with xylose so that the starting inoculum of the *alrA* strain was equivalent to that of the co-culture. CFU/mL of the *alrA* knockdown were not significantly different between the monoculture (dark red) and co-culture (gray) throughout the course of the experiment (*p*-values from each timepoint range from 0.21 to 0.66, student’s unpaired *t*-test). The black line is the total CFU/mL of the parent/*alrA* knockdown co-culture. *n*=3, error bars represent 1 standard error of the mean.

We hypothesized that cells with full knockdown of *alrA* transcription were able to maintain their growth in biofilm co-culture because the parent-RFP cells were providing the necessary D-alanine. To test this hypothesis, we grew monocultures of an *alrA* strain without *gfp* in the genome as on MSgg-xylose with exogenous D-alanine, and found that D-alanine rescued biofilm colony growth (Fig. 4D). D-methionine, an amino acid that can substitute for D-alanine in cell-wall crosslinking [40], also rescued *alrA* growth on MSgg-xylose plates (Fig. 4D), while other D-amino acids that are not incorporated into the cell wall did not rescue colony growth (Fig. S5E,F), suggesting that D-alanine’s specific role in peptidoglycan synthesis is rescued. Thus, sharing of D-alanine within a biofilm rescues *alrA*-depleted cells, likely by stabilizing mutant cell walls.

To test whether *alrA* cells are rescued by the parent-RFP strain in liquid MSgg-xylose, we grew liquid co-cultures and plated dilutions at hourly time points to quantify survival (separating the two strains based on fluorescence). The vast majority of *alrA* cells died within hours in both liquid monocultures and co-cultures with the parent-RFP strain (Fig. 4E). Thus, the rescue of *alrA* knockdown cells by D-alanine sharing in co-cultures is specific to growth in a colony, presumably due to the close proximity of cells that facilitates D-alanine sharing.

### *alrA* knockdown cells stably co-exist with extracellular matrix-deficient wild-type cells

Secretion of extracellular matrix provides structural integrity to colony biofilms [41], including for *B. subtilis* strain 3610 on MSgg agar [42]. Thus, we wondered if matrix plays a role in the rescue of *alrA*, either by providing structural support to *alrA* knockdown cells with weaker walls (Fig. 4C) or by facilitating the diffusion of D-alanine. To test this idea, we deleted the genes encoding both of the main extracellular matrix components (EpsH, TasA) from the parent-RFP strain and the *alrA* knockdown in the *sacA::gfp* wrinkling-proficient library. We mixed the two matrix-deficient strains and quantified colony size and competitive fitness in full knockdown conditions on MSgg-xylose plates. As expected, matrix-deficient co-culture colonies were smaller than matrix-proficient co-cultures, as matrix is necessary for robust biofilm colony growth [11] (Fig. 5A). Nonetheless, matrix-deficient co-cultures exhibited approximately the same fraction of *alrA* cells as matrix-proficient co-cultures (Fig. 5A,B, S6A). Thus, matrix is not required for the growth rescue of *alrA*-knockdown cells.

**Figure 5:**
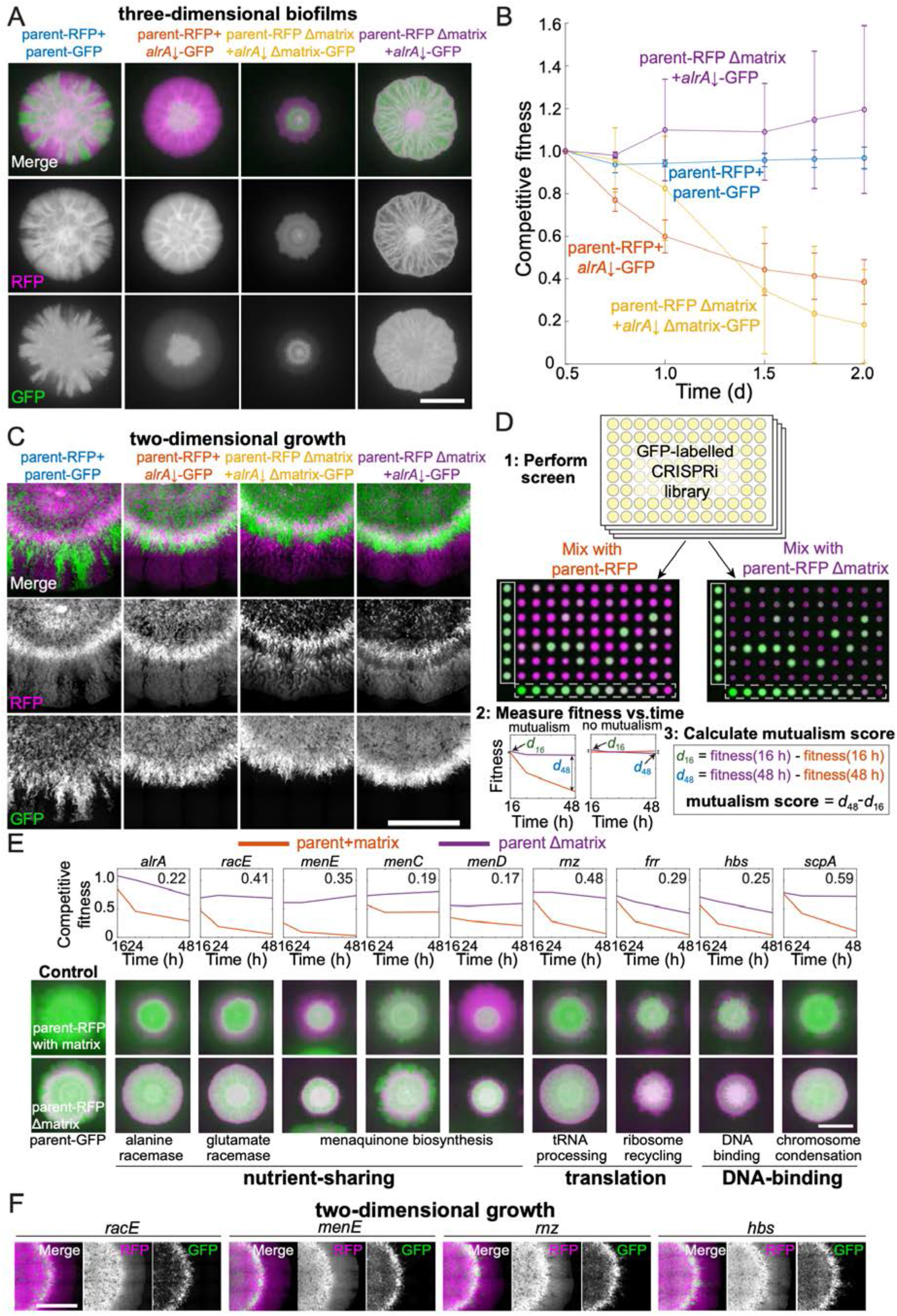
Mutualisms emerge when a nutrient-deficient mutant is the sole provider of extracellular matrix. A) Extracellular matrix is not required for *alrA* rescue (third column), and rescue is enhanced when the *alrA* full knockdown is combined with a matrix-deficient parent strain (fourth column). In the left column, the parent-GFP strain is false-colored in green, and in the other columns the *alrA* knockdown strain is green. The parent-RFP strain is false-colored in magenta. In the first two columns, both strains express matrix proteins. In the third column, both strains lack *epsH* and *tasA*, which encode key matrix components. In the fourth column, *epsH* and *tasA* are deleted from the parent-RFP strain while the *alrA* knockdown produces matrix. Images were acquired after 48 h of growth. Scale bar: 5 mm. B) The competitive fitness of the *alrA* full knockdown decreased over time when both strains or neither produce matrix, while fitness remained 1 (equal proportions of the two strains) and stable over time in co-cultures when the parent is matrix-deficient. The matrix-deficient parent-RFP+*alrA* knockdown (purple) and the parent-RFP+parent-GFP control (blue) data were not significantly different (*p*>0.2 at all time points); nor were the parent-RFP+*alrA* knockdown (orange) and the matrix-deficient parent-RFP+matrix-deficient *alrA* knockdown (yellow) data (*p*>0.07, Student’s *t*-test). The matrix-deficient parent-RFP+*alrA* knockdown (purple) data are significantly higher than the parent-RFP+*alrA* knockdown (orange) at all time points following 0.5 d (*p*<0.014). Data were normalized so that the 0.5 d timepoint is set to 1. Curves are means and error bars represent 1 standard deviation (*n*=2-3 biological replicates). Statistical analysis: Student’s unpaired *t*-test, Benjamini-Hochberg multiple tests correction applied. C) Cells with full knockdown of *alrA* were outcompeted by the parent-RFP strain during two-dimensional growth. Co-cultures were spotted on a MSgg-xylose agar pad and limited to growth in a layer with thickness one cell by applying a cover slip over the cells. Images were acquired after 24 h of growth. Scale bar: 0.4 mm. D) Design of screen to identify mutants that exhibit an increase in fitness when co-cultured with a matrix-deficient parent on MSgg-xylose agar. The *sacA::gfp* library was used in this screen; plate 3 from the 16 h timepoint is shown. The distance between the centers of each colony is 9 mm. White box, controls; white dashed box, titration row. E) Full knockdowns that exhibited mutualism generally involve genes related to nutrient sharing, translation, and DNA-binding. Top: competitive fitness of co-cultures with the wild-type parent-RFP (purple) and matrix production-deficitent parent (red) at 16, 24, and 48 h. Numbers in the top right indicate the mutualism score. Bottom: merged images of biofilm-colony co-cultures on MSgg-xylose agar at 48 h. Scale bar: 5 mm. F) Full knockdowns that exhibited mutualism in three-dimensional biofilms were out-competed by the matrix-deficient parent-RFP strain when growth was limited to two dimensions as in (C). Images were acquired after 24 h of growth. Scale bar: 0.4 mm. For (A,C-F), GFP is false-colored in green and RFP is false-colored in magenta.

Surprisingly, combining the matrix-proficient and wrinkling-proficient *alrA* strain under full knockdown with the matrix-deficient (Δ*epsH* Δ*tasA*) parent-RFP strain resulted in an increased fraction of *alrA*-depleted cells relative to co-cultures with the matrix-proficient parent-RFP strain. In addition to improved growth of the *alrA* knockdown, the RFP fluorescence of the matrix-deficient parent-RFP was also higher and occupied a larger area than in the co-culture of matrix-deficient *alrA* and parent-RFP strains (Fig. 5A). Moreover, rather than a steady decrease in competitive fitness over time (Fig. 5A,B), the competitive fitness of *alrA* against the matrix-deficient parent was stable at ~1 for 48 h. The increased and stable fitness of this strain combination suggests a synthetic mutualistic interaction, defined here as the fitness of both strains in co-culture being stable and higher than either strain on its own, in which matrix-deficient parent-RFP cells restore viability to *alrA* knockdown cells by providing D-alanine, and in turn *alrA* knockdown cells enhance growth of the parent-RFP strain by providing extracellular matrix.

Our finding that the competitive fitness of *alrA* decreased over time in a co-culture with parent-RFP in which both strains were matrix-deficient (and hence did not form a canonical biofim) (Fig. 5A) indicates that the mutualism between *alrA* and the parent-RFP strain requires growth in a matrix-capable biofilm. A recent study showed that biofilm colony expansion in three dimensions depends heavily on extracellular matrix, while two-dimensional growth relies more on cell growth and division [43]. To test whether the mutualism between *alrA* and the matrix-deficient parent was dependent on three-dimensional growth, we grew co-cultures between an agar pad and a coverslip. In this configuration, the growing edge of the colony has a thickness of only one cell, essentially constraining growth to two dimensions (Fig. S6B). Now, in all matrix-production combinations of the *sacA::gfp alrA* strain under full knockdown and parent-RFP, including matrix-proficient *alrA* with the matrix-deficient parent, the parent-RFP strain had a growth advantage at the edge of the colony and out-competed the *alrA* knockdown (Fig. 5C, S6A). Thus, the mutualism we discovered between *alrA* under full knockdown and the matrix-deficient parent is can be eliminated by removing the ability of the colony to grow in three dimensions.

### Many metabolism-related mutants exhibit enhanced rescue in three-dimensional biofilms with matrix-deficient wild-type-like cells

Our discovery that *alrA* knockdown cells grew in a mutualism with an extracellular matrix-deficient parent (Fig. 5A,B) led us to hypothesize that other essential-gene knockdowns might display enhanced growth when cultured with the matrix-deficient parent. To test this hypothesis, we performed co-culture screens of each strain in the *sacA::gfp* library with either the matrix-proficient or matrix-deficient parent-RFP strain on MSgg-xylose (Fig. 5D, S6C, Table S2). We measured competitive fitness at 16, 24, and 48 h to identify strains that had increased and relatively stable fitness when they were the sole provider of extracellular matrix relative to competition with the wild-type-like parent-RFP. For each strain, we calculated a mutualism score defined as the fitness increase due to deletion of matrix components from the parent at 48 h compared with the fitness increase at 16 h; a positive score reflects a growing benefit of being the sole matrix provider, and hence implies relatively stable fitness (Fig. 5D, Fig. S6D). We focused on all strains with a mutualism score 2 standard deviations above the mean across controls (>0.22, Fig. S6D).

On MSgg-xylose, in addition to *alrA*, eight other essential-gene full knockdowns exhibited a high mutualism score (Fig. 5E, S6E,F, Table S5). Of these strains, knockdowns of genes encoding the glutamate racemase (*racE*) and enzymes involved in menaquinone (vitamin K2) synthesis (*menE*) were candidates for nutrient- and matrix-sharing mutualisms similar to that of *alrA*. Knockdowns of two other genes in the menaquinone synthesis pathway (*menC* and *menD*) displayed mutualism scores slightly below 0.22 (Fig. 5E, Table S4), providing further support for a menaquinone-based mutualism. The other essential-gene knockdowns with high mutualism scores encode proteins that bind DNA (*scpA* and *hbs*) or are related to translation (*rnz* and *frr*); these genes likely either play an indirect role in regulating biosynthesis of a shared nutrient to support the mutualism, or employ other mechanisms outside of nutrient sharing. In addition, four non-essential strains exhibited high mutualism scores (Fig. S6F), indicating that the benefits of matrix sharing can extend to genes that are not critical for growth as a monoculture.

Since the *alrA* full knockdown exhibited mutualism in a three-dimensional biofilm but not when growth was confined to two dimensions, we tested whether the other mutualisms were maintained during two-dimensional growth. We grew *racE, menE, rnz*, and *hbs* full depletions in individual co-cultures with the matrix-deficient parent-RFP between a glass slide and a coverslip. In each case, the knockdown was out-competed by the parent-RFP strain at the growing edge of the colony (Fig. 5F). In sum, these data suggest that matrix-dependent mutualisms generally require growth in a three-dimensions.

## Discussion

Here, we created two new libraries of essential gene knockdowns in a biofilm-capable *B. subtilis* strain, and developed a powerful high-throughput screen of competition in bacterial co-cultures to reveal genetic interactions specific to growth in three-dimensional colonies. First, we showed that basal knockdown of some ribosomal proteins reduces competitive fitness with a wild-type-like strain in co-culture colonies (Fig. 2), suggesting a high degree of selection on these genes during colony growth. Second, we found that medium composition can dramatically alter competition (Fig. 3), highlighting the role of the extracellular environment during evolution in a multicellular context. Third, we discovered that knockdown of *alrA* can be rescued through sharing of D-alanine in a three-dimensional biofilm, a context in which the gene is “less essential,” but not in liquid or when growth is confined two dimensions between an agar surface and a coverslip (Fig. 4, 5). Finally, we uncovered a mutualism between *alrA* knockdown cells and a parent deficient in extracellular matrix production based on sharing of nutrients and matrix components, and used this finding to identify several other essential gene knockdowns exhibiting similar mutualistic interactions (Fig. 5). These findings illustrate how growth in a colony/biofilm can alter natural selection by supporting mutant cells that are less likely to survive on their own through short-range interactions.

Despite previous studies showing that D-alanine levels are undetectable in *B. subtilis* 168 liquid culture supernatants [44], we found that D-alanine sharing in a biofilm can rescue *B. subtilis* 3610 mutants that cannot synthesize their own D-alanine. Thus, D-alanine is produced and secreted in a biofilm by wild-type cells at sufficient levels to support growth of the mutant. Since full knockdown of *alrA* causes cells to bulge and die in liquid culture within hours (Fig. 4C,E), we infer that rescue likely occurs early in biofilm development prior to the period of substantial cell death that is thought to drive wrinkling [13], suggesting that rescue is not due to the release of D-alanine by dying cells. The close proximity of cells within the biofilm may aid in rescue, even if secreted D-alanine levels are low. Regardless, this rescue demonstrates that cell-wall synthesis mutants can be supported in native environments through sharing of cell-wall components, which could be provided by many other bacterial species in a multispecies community due to the common chemical makeup of peptidoglycan cell walls [40].

Our observation that rescue of *alrA*-depleted cells did not occur when growth was constrained to two dimensions (Fig. 5C), combined with the finding that rescue occurred in co-culture biofilm colonies when both the *alrA* knockdown and the parent were matrix-deficient (Fig. 5A,B), indicates that some aspect of three-dimensional growth beyond matrix production is fundamental to the rescue. One possibility is liquid uptake facilitated by colony architecture, which has been hypothesized to act like a sponge and thereby drive colony expansion (even more so when extracellular matrix is present) [11, 45]. It is also possible that cellular differentiation and development are disrupted by limiting growth to a thin layer. Irrespective, our findings highlight the importance of future high-throughput genetic screens that embrace the natural context of three-dimensional colony growth on surfaces.

The stable mutualism that we discovered between the *alrA* knockdown and a matrix-deficient parent (Fig. 5B) resembles the initial behavior of Δ*tasA* and Δ*epsH* mutants grown together as pellicle biofilms on liquid surfaces, in which the pellicle architecture is preserved by cross-complementation for many passages [46]. Our discovery of matrix-based mutualisms involving multiple genes with a range of cellular functions (Fig. 5E) motivates future studies to probe the nature of the *tasA-espH* interaction in these interactions, specifically to examine which matrix components are most important and whether mutualisms can be sustained through repeated passages or with different starting ratios of the strains in the co-culture. Importantly, the fact that these mutualisms appear to generally require growth in three dimensions further highlights the importance of the three-dimensional geometry of the native environment during evolution.

Together, our results demonstrate that growth in a biofilm can drive genetic diversity and illustrate the potential for mutualism between nutrient and matrix sharing in native biofilms. Such mutualisms may occur during plant root colonization, when the bacterial extracellular matrix is particularly important and may serve to pull nutrients from the root and surrounding soil [8]. In addition to the potential implications for plant growth-promoting bacteria in the rhizosphere, this study provides a foundation to understand how microbial biofilm growth affects selection in industrial and clinical settings.

## Supporting information

Supplemental Table 1 - strains list

Supplemental Table 2 - library data values

## Acknowledgments

The authors thank the Huang lab and Petra Levin for helpful discussions, Nicola Stanley-Wall for providing the *sacA::gfp* construct, and Dan Kearns for providing matrix mutant constructs. The authors acknowledge support from the Allen Discovery Center at Stanford on Systems Modeling of Infection (to H.A.A. and K.C.H.), the Stanford Bioengineering Summer Research Experience for Undergraduates Program (to H.G.), and NIH K22 Award AI137122 (to J.P.). K.C.H. is a Chan Zuckerberg Biohub Investigator.

## METHODS

### Media

Strains were grown in LB (Lennox broth with 10 g/L tryptone, 5 g/L NaCl, and 5 g/L yeast extract) or MSgg medium (5 mM potassium phosphate buffer, diluted from 0.5 M stock with 2.72 g K_2_HPO_4_ and 1.275 g KH_2_PO_4_, and brought to pH 7.0 in 50 mL; 100 mM MOPS buffer, pH 7.0, adjusted with NaOH; 2 mM MgCl_2_•6H_2_O; 700 μM CaCl_2_•2H_2_O; 100 μM FeCl_3_•6H_2_O; 50 μM MnCl_2_•4H_2_O; 1 μM ZnCl_2_; 2 μM thiamine HCl; 0.5% (v/v) glycerol; and 0.5% (w/v) monosodium glutamate). MSgg medium was made fresh from stocks the day of each experiment for liquid cultures, or a day before the experiment for agar plates. Glutamate and FeCl_3_ stocks were made fresh weekly. Colonies were grown on 1.5% agar plates. For nutrient addition assays (Fig. 3, S4), we supplemented LB with one of the following: 0.5% (w/v) monosodium glutamate, 2 mM MgCl_2_•6H_2_O, 50 μM MnCl_2_•4H_2_O, 2 mM MgCl_2_•6H_2_O, 0.5% (w/v) L-asparagine, 0.5% (w/v) L-aspartic acid, 0.5% (w/v) L-lysine, or 0.5% (w/v) D-glutamic acid, and we supplemented MSgg with one of the following: 0.5% (w/v) L-cysteine, 0.5% (w/v) L-glutamine, 0.5% (w/v) L-glycine, 0.5% (w/v) L-serine, or 0.5% (w/v) L-tryptophan. Where indicated, L-threonine was added to MSgg at a concentration of 0.1 mg/mL. D-amino acids (D-alanine, D-methionine, D-glutamate, D-leucine, D-serine, D-valine) were each used at a concentration of 0.04 mg/mL. We made TY medium for phage transduction using the LB recipe above supplemented with 10 mM MgSO_4_ and 0.1 mM MnSO_4_. Antibiotics for selection of mutant strains were used as follows: kanamycin (kan, 5 μg/mL), MLS (a combination of erythromycin at 0.5 μg/mL and lincomycin at 12.5 μg/mL), chloramphenicol (cm, 5 μg/mL), tetracycline (tet, 12.5 μg/mL) and spectinomycin (spc, 100 μg/mL).

### Strain construction

All strains and their genotypes are listed in Table S1. For library construction, we used SPP1 phage transduction [47]. We used a 168 strain containing *P_xyl_-dCas9* at the *lacA* locus (CAG74399) as a donor and wild-type strain 3610 (a gift from Dan Kearns) as the recipient to create the 3610-dCas9 parent strain (CAG74331) using MLS for selection. We then used this 3610-dCas9 parent as the recipient and a strain with *P_spachy_-gfp* at the *sacA* locus (NRS1473, gift from Nicola Stanley Wall) or a strain with *P_veg_-gfp* at the *thrC* locus (HA47, construction described below) as the donor strain to create the parent-GFP strain expressing *dCas9* and *gfp*, using kanamycin and tetracycline for selection, respectively.

For construction of mutant strains, we used either the *sacA::gfp* parent or the *thrC::gfp* parent as the recipient strain and strains from a 168 CRISPRi library [30] as the donor. We amended the phage transduction protocol to increase the throughput of strain construction as follows. We grew donor strains in 96-well deep-well plates (1-mL cultures in TY medium in 2-mL wells) for at least 5 h shaking at 37 °C with a Breath-easy (Sigma-Aldrich) film covering the plate. We then aliquoted 0.1 mL of 10^−5^ dilutions of fresh phage stocks grown on strain 3610 cells (10^−5^ was chosen as the dilution factor because it provided the appropriate level of lysis for our phage stock in a trial transduction) into 77 or 71 glass test tubes (each plate of the library contains 77 strains, except the fourth plate contains 71 strains). We added 0.2 mL of each culture to the tubes and incubated the entire rack at 37 °C for 15 min. Then, working quickly in batches of 11, we added 4 mL of TY molten soft agar (~55 °C) to each phage-cells mixture, mixed gently, and poured onto TY plates so that the soft agar covered the entire plate. We incubated these plates at 37 °C overnight in a single layer (not stacked). The next day, we examined the plates for lysis and added 5 mL TY broth with 250 ng DNase to each plate and scraped the top agar with a 1-mL filter tip to liberate phage. We then pipetted the TY broth into a syringe attached to a 0.45-μm filter and carefully filtered into a 5-mL conical vial. After filtering, 1 mL of lysate was added to the appropriate well of a deep-well 96-well plate. Once all of the phage was isolated, we arrayed 10 μL of each phage stock into 96-well microtiter plates. We aliquoted 100 μL of a saturated (>5 h of culturing, OD_600_>1.5) culture into the wells containing phage and incubated for 25 min at 37 °C without shaking. We plated the phage/cell mixtures onto selection plates (LB with chloramphenicol and citrate to select for the sgRNA locus) and incubated the plates for 18 h at 37 °C. Any plates that did not have visible colonies after this incubation were incubated further at room temperature, and colonies generally appeared within a day. We streaked transductant colonies for single colonies onto LB+chloramphenicol plates and stocked a single colony for each strain in the library by growing in 5 mL LB on a roller drum at 37 °C to mid- to late-log phase and then adding the culture to the appropriate well of 96-well plate with a final concentration of 15% glycerol. The library was stocked at −80 °C.

The GFP-labeled Δ*epsH* Δ*tasA alrA* knockdown strain (HA823) and the parent-RFP Δ*epsH* Δ*tasA* strain (HA825) were constructed using phage transduction as described above, using DS9259 and DS3323 (gifts from Dan Kearns) as donor strains for *epsH::tet* and *tasA::tn10spc*, respectively, and the *sacA::gfp alrA* knockdown strain (HA761) or parent-RFP (HA12) as the recipient. The *epsH::tet* transduction was performed first, and the resulting strains were used as the parent to add the *tasA::tn10spc* construct.

To construct plasmid pDG1731-gfp (*P_veg_-sfGFP* in a *thrC* integration construct), the following primers were used to clone superfolder GFP (sfGFP) and add the *P_veg_* promoter: forward, tcctagaagcttatcgaattcCTTATTAACGTTGATATAATTTAAATTTTATTTGACAAAAATGGGCTCGTGTTGTACAATAAATGTAACTACTAGTACATAAGGAGGAACTACTATGAGCAAAGGAGAAGAACTTTTC; reverse, ttaagcaccggtttattaTTTGTAGAGCTCATCCATGCC. The amplicon and pDG1731 were both digested with HindIII and AgeI and ligated together. The ligation was used to transform chemically competent *E. coli*. We transformed *B. subtilis* 168 with pDG1731-gfp to create HA45 and confirmed double crossover (spc^R^, MLS^S^), then used HA45 as the donor and HA2 as the recipient in phage transduction to create the *thrC* parent-GFP strain (*P_xyl_-dCas9 thrC::P_veg_-gfp*, HA47).

### Growth conditions for library screens of growth on agar plates

To grow the library for monocultures and co-cultures, we inserted a sterile 96-well Singer pin (Singer Instruments) into frozen glycerol stocks and applied pressure and agitation so that each pin picked up some of the frozen glycerol stock from the appropriate well. The Singer pin was used to spot onto LB agar in a rectangular Singer plus plate and the plate was incubated overnight at 37 °C. A sterile 96-well Singer pin was used to pick up cells from each colony and inoculate 200 μL of LB in a 96-well plate. The parent-GFP strain was inoculated in some of the empty wells on the edge of each plate as controls. The plate was covered with an AeraSeal breathable film (Sigma-Aldrich), and grown on a plate shaker at 37 °C for 4-5 h until all wells were cloudy (OD_600_~1.0).

One hundred microliters of each culture were pipetted into a separate 96-well plate with 100 μL of a parent-RFP culture in each well. This plate was used as the inoculum for the competitive fitness screen. The remainder of the cultures in the original 96-well plate were used as the inoculum for the monoculture screen. To quantify competitive fitness, a titration row of parent-RFP and parent-GFP mixtures in 10% increments (100% parent-RFP+0% parent-GFP; 90% parent-RFP+10% parent-GFP, etc.) was added to each plate (Fig. 1D). Since the oxygen limitation that results in stationary cultures causes cell death in *B. subtilis* [21], we ensured that the library was aliquoted and spotted within 1 h. For most assays, a Singer ROTOR HDA pinning robot (Singer Instruments) was used to pin ~1 μL of liquid cultures onto LB or MSgg agar Singer plus plates (with 35 mL of medium poured on a level surface for co-cultures, or 50 mL of medium for monocultures), without and with xylose. We used the “spot many” protocol of the Singer ROTOR HDA to mix the wells before spotting and transferred 12 times from the source liquid plate to the target agar plate. For some assays, a RAININ Benchsmart 96-well pipetting robot was used rather than the Singer ROTOR HDA to pipet 1 μL onto the agar plates. Agar plates were incubated at 30 °C and placed in a box or were loosely covered in plastic to reduce drying.

When screen outliers (Fig. 2D, 3C) were replicated, strains were streaked for single colonies, which were inoculated into 200 μL of medium in the interior wells of a 96-well microtiter plate. The exterior wells were inoculated with the parent-GFP control strain, leaving the top for the parent-GFP+parent-RFP titration. To replicate findings regarding *alrA*, fresh colonies of the *alrA* knockdown strain and the parent-RFP strain were inoculated into 5 mL LB in test tubes and cultured on a roller drum at 37 °C to an OD_600_~1.0. Equal-volume mixtures of the cultures were spotted along with a parent-GFP+parent-RFP titration in 12- or 6-well plates.

### Imaging and image analysis of monoculture colonies in the library

Monoculture colonies were imaged using a Canon EOS Rebel T5i EF-S with a Canon Ef-S 60 mm f/2.8 Macro USM fixed lens. The DSLR camera was set up at a fixed height in a light box with diffuse lighting from three sides. The lighting and camera settings were maintained for the duration of the experiment, using the “manual” mode on the camera. The EOS Utility software was used to run the camera. Plates were imaged colony side up to avoid imaging through the agar. Images were analyzed using FIJI and scored as “grew outside original spot”, “did not grow outside original spot”, or “died and/or threw off suppressors.” The ones classified “died and/or threw off suppressors” were assigned a colony size of 0. Suppressors were identified based on off-center colonies, often in flower petal-like arrangements in which one or a few cells within the original spot eventually grew but the majority of the cells did not. Colony size was measured manually in FIJI by drawing a diagonal line across the diameter of the colony.

### Imaging and image analysis of biofilm co-cultures

A Typhoon™ FLA 9500 scanner was used to image colonies using the multi-plate drawer. We used ~35 mL of medium with agar on a Singer rectangular plate to be near the plane of focus when imaging through the agar. GFP (473 nm laser, Long Pass Blue filter) and RFP (532 nm laser, Long Pass Green filter) signals were acquired.

For image analysis, Typhoon RFP and GFP images were cropped to contain only one plate and the image was rotated so that A1 was in the top left corner. Custom Matlab code was written to read in each plate, divide it into a grid in which each grid cell contained one colony, and extract the fluorescence level across that grid. The ratio of the extracted GFP and RFP values was computed for every colony, and the ratio values for the titration row against the fraction GFP was fitted using the function *I* = *aG*/(1-*bG*), where *I* is the GFP/RFP ratio, *G* is the fraction GFP, and *a* and *b* are fit parameters (Fig. 1D). The fit parameters from the titration row were used to map the library data and assign assign GFP fractions; the data were normalized so that the average of the internal parent-RFP and parent-GFP co-culture controls was 1.

### CRISPRi *rfp* knockdown

Wild-type 3610, the parent-RFP, and the CRISPRi-RFP strains were cultured in 5 mL test tubes at 37 °C to an OD_600_~1 in liquid LB. The parent-RFP strain was spotted onto LB and MSgg agar plates without xylose, while the CRISPRi-RFP strain was spotted onto LB and MSgg agar in 12-well plates containing 0.0005% to 1% xylose. The RFP fluorescence of the colonies was imaged as described above, and FIJI was used to quantify the fluorescence intensity of each colony, using wild-type 3610 as a blank.

### Wild-type 3610, parent-GFP, and parent-RFP biofilm and non-biofilm colony growth

Wild-type 3610, the parent-GFP, and the parent-RFP strains were cultured in liquid LB to an OD_600_~1 at 37 °C. One microliter of each culture was spotted onto LB or MSgg agar in a 6-well plate. Colonies were cultured for 48 h at 30 °C in a plastic bag with a wet paper towel to increase humidity. Colonies were imaged using the DSLR setup as described above.

### Liquid culture for growth rate analysis

A single colony was used to inoculate 200 μL LB in a 96-well microtiter plate. OD_600_ was measured every 7.5 min using a Biotek Epoch plate reader at 37 °C. OD_600_ curves were blanked and smoothed. The maximum growth rate of each culture was defined as the maximum derivative of ln(OD_600_).

### Model of nutrient-dependent colony growth

To determine how competitive fitness in a co-culture colony is affected by differences in growth rate, we simulated a reaction-diffusion model in which two cell types with densities (*C_i_*, *i* = 1,2) are inoculated in a circular spot from which they spread randomly in two dimensions to compete for fresh nutrients (*n*) and grow with distinct maximal growth rates (*M_i_*), according to the following equations:

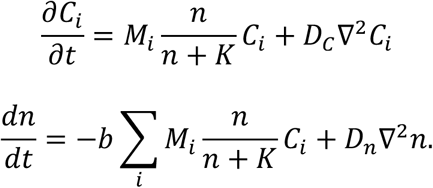

*D_c_* is the cell diffusivity, *D_n_* is the diffusivity of nutrients, *b* is a conversion factor dictating how nutrients lead to cell growth, and cell growth is limited by nutrients when *n~K* or less. Initially *n* = *n*_0_ everywhere and, to represent the initial pinning of cells to the agar surface, *C* = *C_0_* within a disc of radius *r*_0_ and outside of this disc *C* = 0.

The transformations 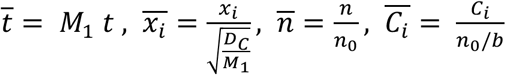 render the equations dimensionless:

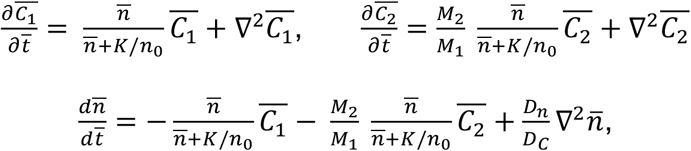

where 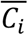 is the ratio of *C_i_* compared with its value if cells of type *i* consume all available nutrients, and 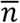 is the ratio of nutrient compared with its initial value (so 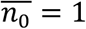). So, the governing dimensionless parameters are 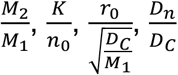 and 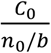.

Several parameters were estimated from data. The maximal growth rate *M*_1_ =0.0175 min^−1^ set the timescale and corresponds to a 40 min doubling time, similar to *B. subtilis* 3610 at 30 °C. The radius of the initial spot (1 mm) set the spatial scale. The ratio of maximal growth rates 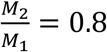 was set to match the ratio in LB for ribosomal knockdowns compared with wild type (Fig. 2C). We estimated the initial areal density of cells compared with their saturation density to be 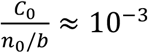 within the initial spot. To obtain this estimate, we assumed 1 μL of stationary phase culture spotted 10^6^ cells over *π* mm^2^, and that the spotted cells saturate at a density of 10^9^ cells/mm^2^ (assuming 1 mL of stationary phase culture contains 10^9^ cells [48] and concentrates into 1 mm^3^ when pelleted, the latter indicating a maximal density of 10^9^ cells/mm^3^ within a colony with height 1 mm). We assumed nutrients diffuse much faster than cells such that *D_n_/D_c_* ≈ 100.

The model approximately recapitulates the competitive fitness data (defined in the model as the ratio of integrated cell densities ∫ *C*_2_ *dA*/ ∫ *C*_1_ *dA*) and colony radius at 16 h when *D_c_* ≈ 0.003 mm^2^h^−1^ (Fig. 2E,inset). Results are fairly robust to variation in 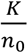 and 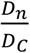 (Fig. S3C).

### DAVID functional enrichment

We used the DAVID functional annotation tool (https://david.ncifcrf.gov) to determine whether particular gene classes were enriched for each phenotype. The BSU identification number of the strains identified by our analysis and of the entire CRISPRi library were used as the “list” and the “background,” respectively, using the “locus tag” option on the website)

### D-amino acid rescue experiments

The unlabelled *alrA* knockdown (HA420) and wild-type 3610 were grown to an OD6_00_~1 in liquid LB. One microliter of each culture was spotted onto MSgg or MSgg-xylose agar plates with 0.04 mg/mL of one of the D-amino acids. Cultures were incubated for 48 h, imaging at 24 and 48 h using the DSLR setup described above.

### Liquid growth of *alrA* monocultures and co-cultures for plating efficiency

Cultures of the *alrA* knockdown strain (HA420) and the parent strain (HA2) were separately cultured from a colony in liquid LB at 37 °C until both strains reached OD_600_ ~1.0. The HA420 and HA2 cultures were mixed 1:1, and the mixture along with the HA420 monoculture were back-diluted 1:200 into 3 mL MSgg medium with 1% xylose and incubated at 30 °C, shaking at an angle. At 0 h, 1 h, 2 h, 3 h, 4 h, and 5 h, cultures were sampled, ten-fold serially diluted, and spotted onto MLS or chloramphenicol selection plates to determine CFU/mL of each strain (HA2 and HA420 are MLS^R^, HA420 is cm^R^, HA2 is cm^S^). We incubated the dilutions overnight and counted colonies to calculate CFU/mL.

### Liquid growth of wild-type 3610 and *alrA* knockdowns for microscopy

The 3610 wild-type strain and an unlabelled *alrA* knockdown strain (HA420) were grown to an OD_600_~1 in LB. Each strain was back-diluted 1:200 into 3 mL MSgg+1% xylose to fully knock down *alrA* transcription during incubation at 30 °C, shaking at an angle. At 0 and 6 h, 1 μL of each culture were spotted onto 1X PBS pads made with 1.5% agar. When dry, we added a coverslip and imaged the cells in phase contrast on a Nikon Ti-E inverted microscope using a 100X objective (NA: 1.4).

### Two-dimensional culturing

Strains were grown in LB to OD_600_~1. While strains grew, we prepared a large agar pad at least an hour before imaging using the bottom of a rectangular Singer PlusPlate culture plate and 30 mL of MSgg+1% xylose. After the agar solidified, we added a second Singer PlusPlate on top to prevent contamination and drying. We mixed the strains 1:1 volumetrically and spotted 0.5 μL of this mixture onto the agar pad. After the spot dried, we added a large 113 by 77 mm custom-made no. 1.5 glass coverslip (Nexterion). The pads were incubated in a plastic bag with a wet paper towel to maintain humidity at 30 °C for 24 h. The entire spot was captured in a grid of images using a Nikon Ti-E inverted microscope with a 40X air objective (NA: 0.95) integrated with μManager [49]. Images were stitched together using custom Matlab (MathWorks) code. GFP and RFP stitched images were merged using Adobe Photoshop, adjusting each channel equally.

### Mutualism screen

The mutualism screen (Fig. 5D) was performed as described above, except two screens were performed side by side: one in which each strain in the *sacA::gfp* library was co-cultured with the parent-RFP strain (HA12), and one in which each strain was co-cultured with the Δ*epsH* Δ*tasA* parent-RFP strain (HA825). These screens were carried out on MSgg+1% xylose plates with a titration row of HA773 (parent-GFP) in combination with either HA12 or HA825, as described above.

### Qualitative fitness determination via dilution streaking

Strains were inoculated from a fresh colony into 5 mL LB and incubated at 37 °C for ~5 h on a roller drum. Cultures were streaked onto agar plates using sterile wooden sticks. A new sterile stick was used for each streak. These plates were incubated overnight (~18 h) at 37 °C, and imaged using the DSLR camera setup described above.

### Statistical Methods

All statistical tests are stated in the figure legends. To determine whether data was significantly different, Student’s unpaired *t*-tests were applied. The Benjamini-Hochberg multiple tests correction was applied to data in Figures 2C, 3C, and 5B.

## Supplemental Figures

**Figure S1:**
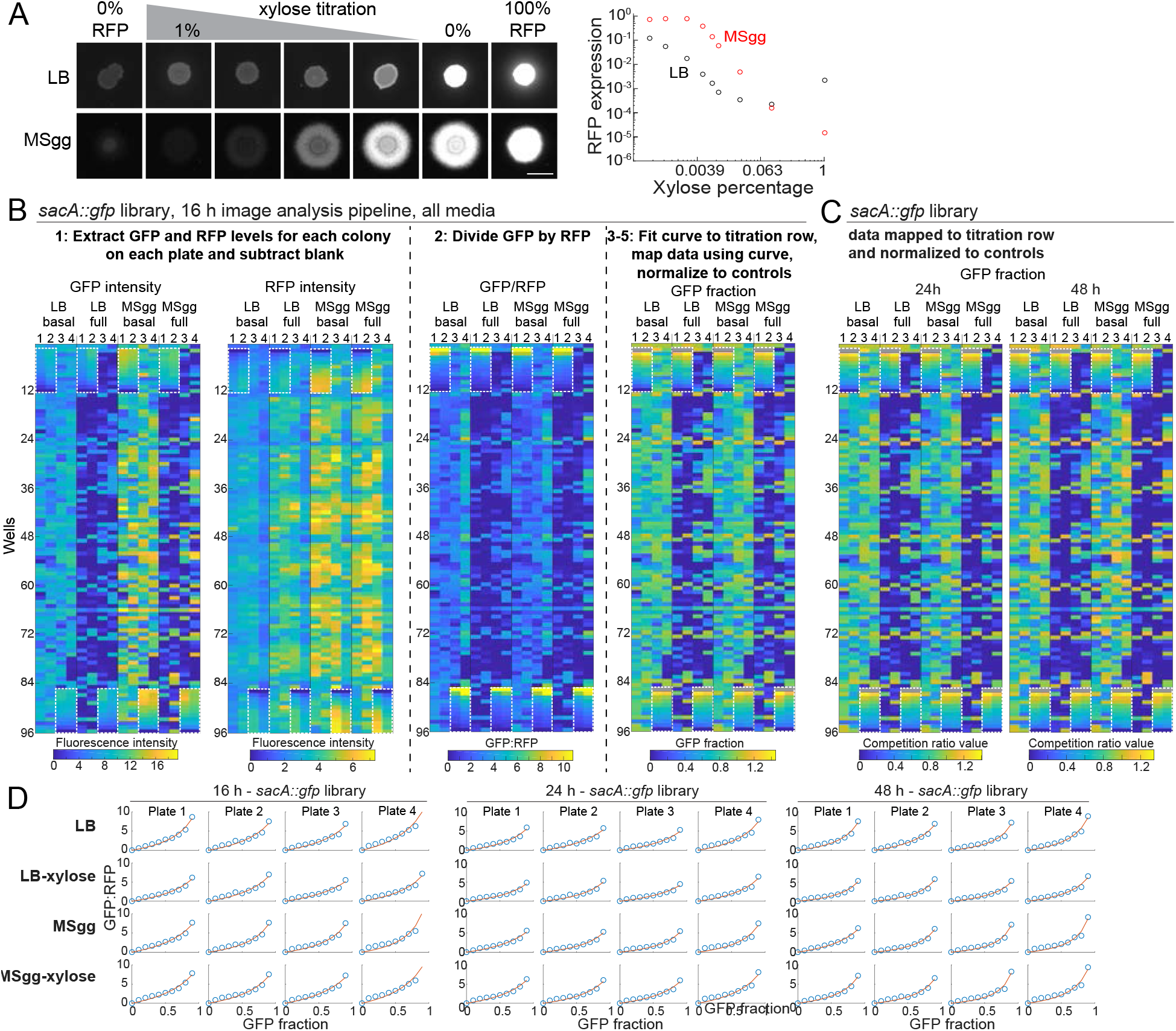
CRISPRi is an efficient tool for tunable knockdown of gene expression in non-biofilm and biofilm colonies, enabling high-throughput competition screens. A) Varying CRISPRi induction generates titrated gene expression in colonies on LB and on the biofilm-promoting medium MSgg. We spotted a strain with CRISPRi targeting *rfp* onto LB or MSgg agar plates with various amounts of xylose xylose. Left: images were acquired after 24 h of growth. Right: RFP levels varied inversely with xylose concentration, with basal repression minimally decreasing expression of RFP and higher levels of xylose repressing expression by 10- to 1000-fold in LB and ~10,000-fold in MSgg. Data were normalized to RFP levels in a strain without a CRISPRi sgRNA (100% RFP). Scale bar: 5 mm. B) Competition data for the entire *sacA::gfp* library after 16 h of growth at each step of the analysis pipeline. The titration row is denoted by white dashed boxes. The few gray boxes represent empty wells or wells that involved division by zero during processing and hence were ignored. C) Competition data for the *sacA::GFP* library after 24 h and 48 h of growth. The titration row is denoted by white dashed boxes. The few gray boxes represent empty wells or wells that involved division by zero during processing and hence were ignored. D) Data from the titration row of parent-GFP and parent-RFP co-cultures were well fit by the predicted equation *I=aG*/(1-*bG*) (red lines, Fig. 1D). Blue open circles show the ratio of GFP:RFP intensities of the 0-90% GFP (100-10% RFP) colonies plotted against the fraction of GFP for each plate of each library at each time point in (B,C).

**Figure S2:**
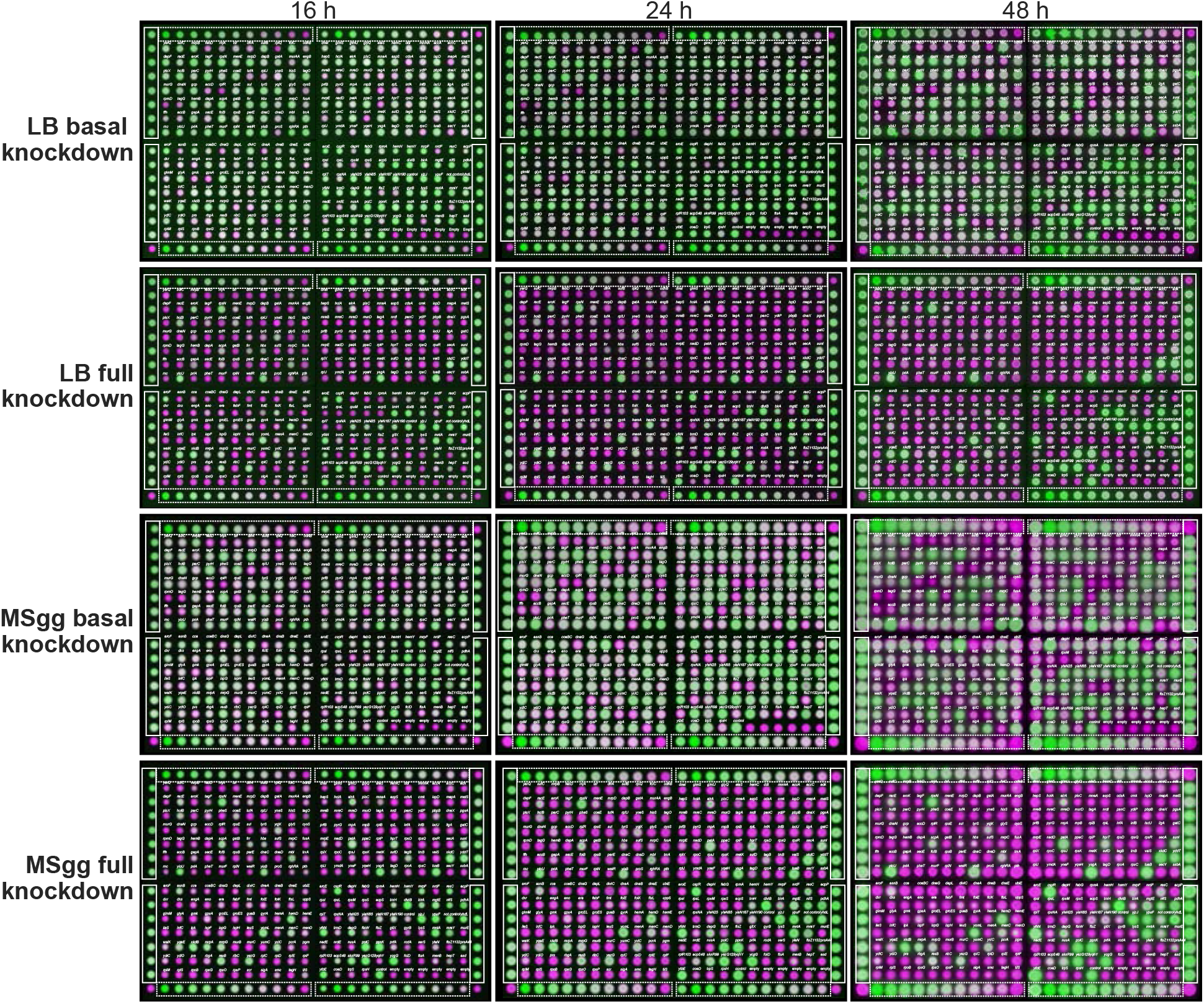
Images of plates from the competition screen with the *sacA::GFP* library. Merged images from the competition screen on LB and MSgg agar under basal and full knockdown, at 16, 24, and 48 h. The CRISPRi strains and parent-GFP controls are false-colored in green and the parent-RFP is false-colored in magenta. The dashed boxes show the titration row of each plate and the solid boxes show the parent-GFP + parent-RFP controls. The distance between the centers of neighboring colonies is 9 mm.

**Figure S3:**
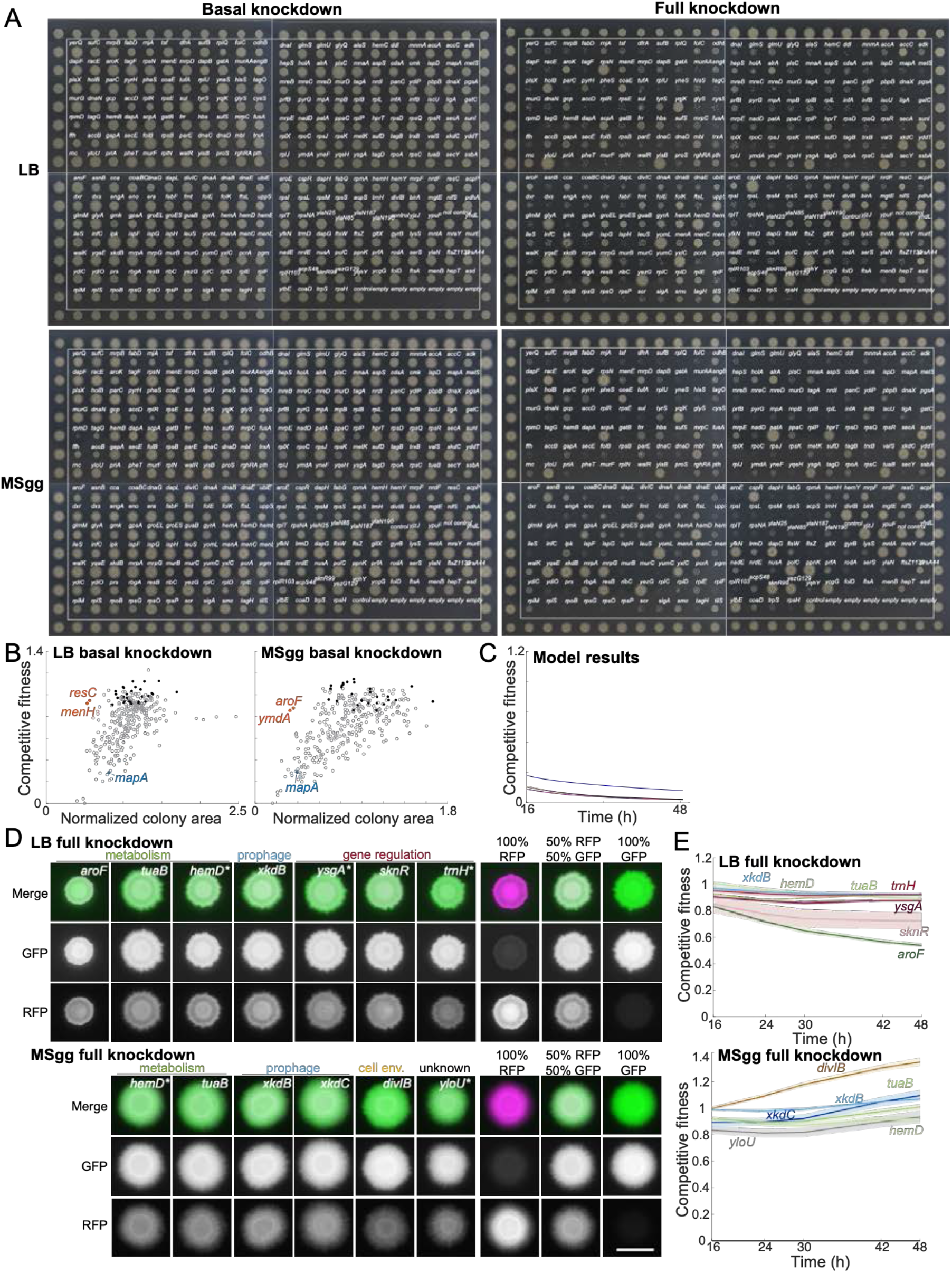
Monoculture colony size screen, and analysis of knockdowns with low and high competitive fitness. A) Monoculture colonies on MSgg agar (basal knockdown) exhibited more variation in size than monoculture colonies on LB agar after 16 h of growth. With full knockdown, there were many small (relative to controls) monoculture colonies on LB and on MSgg after 16 h. The CRISPRi library is within the white boxes and colonies outside of the boxes are parent-GFP controls. The distance between the centers of neighboring colonies is 9 mm. Data for colony areas is in Table S2. B) A few knockdowns exhibited high competitive fitness in co-culture despite having reduced colony sizes in monoculture in basal knockdown conditions (orange), and one non-essential gene knockdown in the library had reduced competitive fitness (blue). The rest of the library are in gray and the controls are in black. C) The reaction-diffusion model of colony growth (Methods) recapitulates competitive fitness when *D_C_* = 0.003 mm^2^ h^−1^ and *M*_1_/*M*_2_ = 0.8; competitive fitness is insensitive to changes in *K*/*n*_0_ and *D_n_*/*D_C_*. Parameters *M*_1_ = 0.0175 min^−1^, *r*_0_ = 1 mm, and *C*_0_/(*n*_0_/*b*) = 0.001 are estimates from data. The black line shows the simulated competitive fitness corresponding to the colony in the inset of Fig. 2E in which both competitive fitness and colony size data were recapitulated with *K*/*n*_0_ = 0.05 and *D_n_*/*D_C_* =100. Competitive fitness was largely unchanged if *D_n_*/*D_C_* = 1000 (dark pink, partly beneath black line) or *D_n_*/*D_C_* = 10 (light pink, partly beneath black line), or if *K*/*n*_0_ = 1.0 (dark blue) or *K*/*n*_0_ = 0.05 (light blue, partly beneath black line). D) Strains that competed well at full knockdown included genes related to metabolism, gene regulation, prophage, and cell envelope (*divIB*), along with *yloU*, a gene of unknown function. Merged images show the parent-RFP false-colored in magenta and the knockdown strain false-colored in green. The 100% parent-RFP, 50% parent-RFP+50% parent-GFP, and 100% parent-GFP controls are shown to the right of the knockdowns. Scale bar: 5 mm. E) Competitive fitnesses of the knockdowns with the highest fitnesses at 16 h were mostly stable over time with full CRISPRi induction. Curves are means and shaded regions represent 1 standard error of the mean (*n*=3 biological replicates).

**Figure S4:**
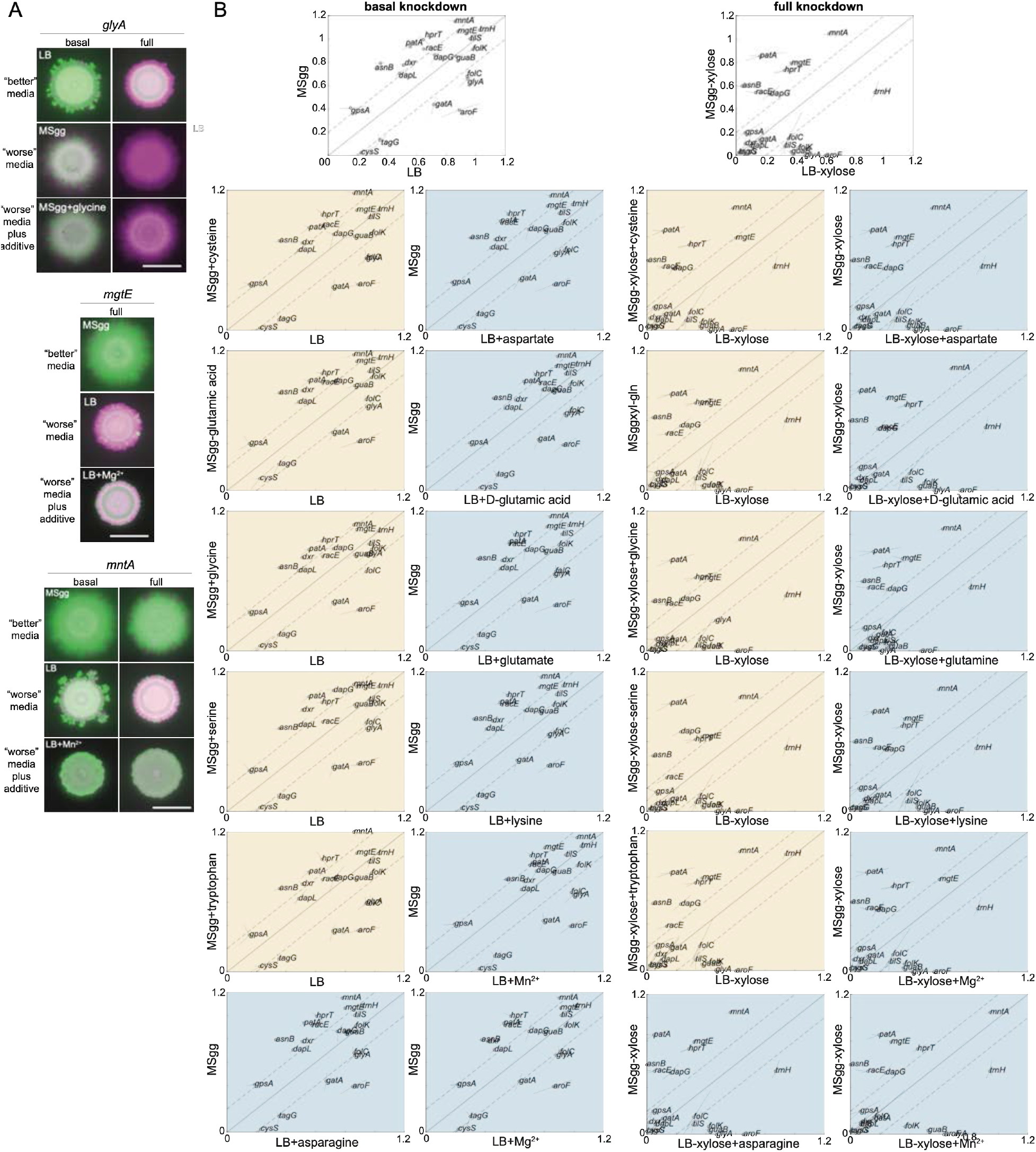
Many nutrients do not impact the competitive fitness of knockdowns with different phenotypes between MSgg agar and LB agar. A) The competitive fitness of *glyA, mgtE*, and *mntA* knockdowns improved when glycine was added to MSgg agar, Mg^2+^ was added to LB agar, or Mn^2+^ was added to MSgg agar, respectively. Images are merges of fluorescence from the knockdown (false-colored in green) and the parent-RFP (false-colored in magenta) after 48 h. Scale bar: 5 mm. B) Many nutrients did not alter competitive fitness. Baseline competitive fitness values (top row) versus when a nutrient was added to MSgg agar (yellow) or LB agar (blue). Means are displayed as black circles, with each replicate at the end of the lines extending from the circle. Data are from the 48 h time point.

**Figure S5:**
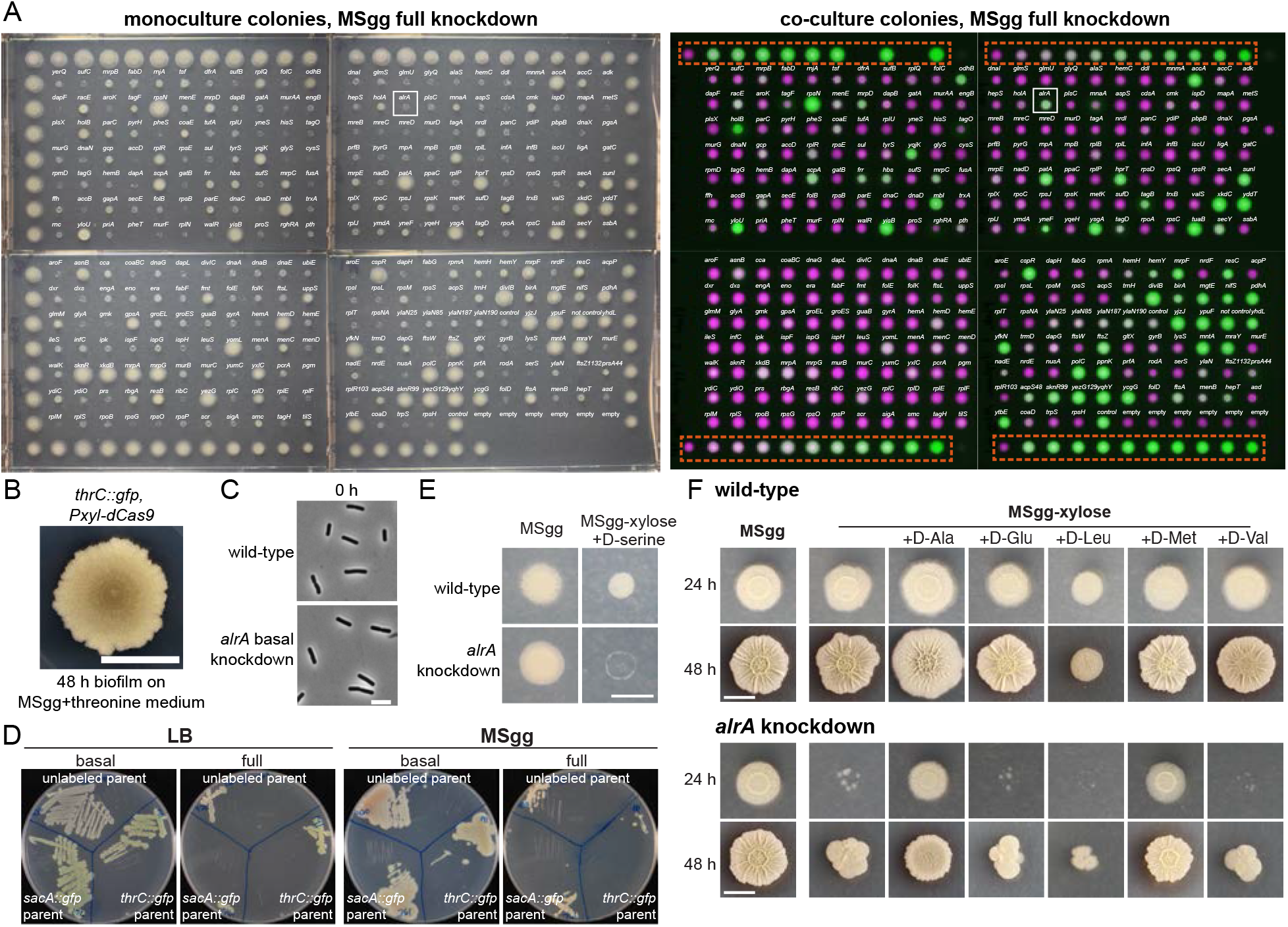
Under full induction, the *alrA* knockdown dies as a monoculture colony but grows when co-cultured with the wildtype-like parent. A) A screen under full CRISPRi induction for knockdowns that die as monoculture colonies (left) but survive in co-culture (right) highlighted *alrA* (white box). Left: monoculture colonies of the *thrC::gfp* library on MSgg-xylose; outer colonies are *thrC::gfp* parent controls. Right: merged images of co-cultures of the *thrC::gfp* CRISPRi library (false-colored in green) with the parent-RFP strain (false-colored in magenta). The titration row is indicated by the red dashed box. The distance between the centers of two neighboring colonies is 9 mm. B) The *thrC::gfp* parent strain did not form wrinkles. One microliter of an LB liquid culture (OD_600_~1) was spotted onto an MSgg-threonine plate and incubated at 30 °C for 48 h. Scale bar: 5 mm. C) Cells with basal knockdown of *alrA* are rod-shaped, similar to wild type. Images were taken directly before adding xylose to fully deplete cells of *alrA* for 6 h, as shown in Figure 4C. Scale bar: 5 μm. D) Full knockdown of *alrA* resulted in a growth defect. Cultures were grown and streaked out onto LB and MSgg plates with and without xylose to qualitatively observe growth under basal and full knockdown. A standard (100-mm) cell culture dish is shown. E) D-serine inhibited the growth of wild-type colonies and did not rescue growth of the *alrA* knockdown. The colony was imaged through agar (to avoid having the objective contact the colony) after 24 h. D-serine was supplemented at 0.04 mg/mL. Scale bar: 5 mm. F) Most D-amino acids did not restore the growth of *alrA*-depleted cells. D-Leu inhibited the growth of wild-type colonies (top). Only D-alanine and D-methionine restored growth of full *alrA* knockdown (bottom). D-amino acids were supplemented at 0.04 mg/mL. We did not test D-Ile, D-Phe, D-Thr, or D-Tyr. Scale bar: 5 mm.

**Figure S6:**
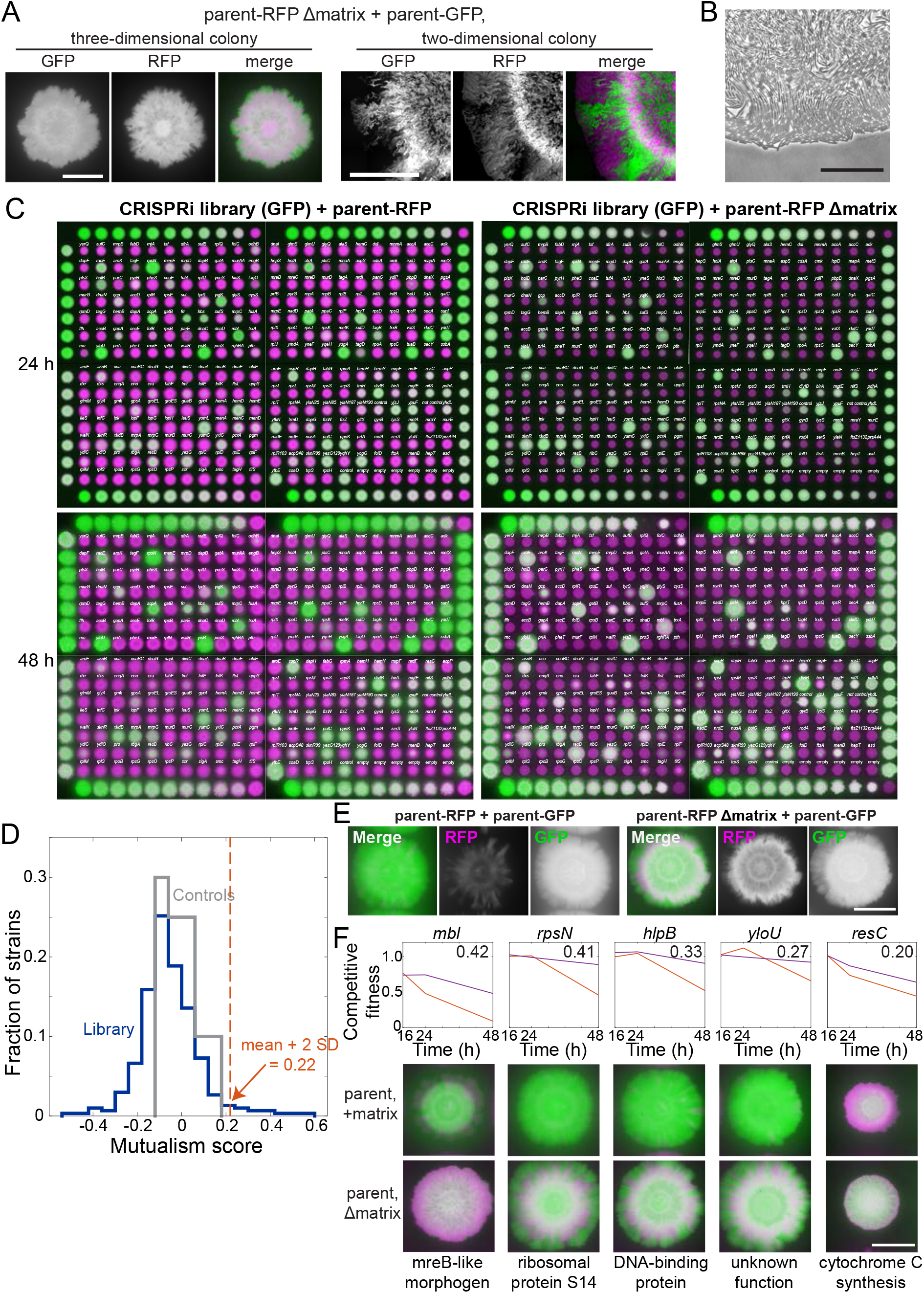
A mutualism screen reveals full knockdowns with improved growth in co-culture when the parent is deficient in production of extracellular matrix. A) The matrix-deficient parent-RFP+matrix-proficient parent-GFP co-culture did not form sectors in a three-dimensional colony (left), but did at the edge of a two-dimensional colony (right). Merges show the parent-GFP and parent-RFP strains false-colored in green and magenta, respectively. Left scale bar: 5 mm; right scale bar: 0.4 mm. B) The leading edge of a co-culture grown between agar and a coverslip is one cell thick. One microliter of cell culture was spotted onto an MSgg-agar pad and a coverslip was applied to limit growth to two dimensions. The culture was incubated for 24 h and the colony edge was imaged. Scale bar: 50 μm. C) Results from a mutualism screen comparing the competitive fitness of knockdown strains co-cultured with a matrix-proficient (left) or matrix-deficient (right) parent. Control parent-RFP+parent-GFP co-cultures are located on the right and left edges of the library, and the titration row is shown on the top and bottom rows. The distance between the centers of neighboring colonies is 9 mm. GFP (from parent-GFP or knockdown strains) and RFP (from parent-RFP) fluorescence signals are false-colored in green and magenta, respectively. D) The library exhibited a wide range of mutualism scores, with 11 full knockdowns exhibiting a mutualism score >2 standard deviations higher than the across all strains (>0.22). The library is shown in blue and the controls are shown in gray. E) Representative controls of the parent-RFP strain grown with the parent-GFP strain showing the final composition of RFP and GFP in the colonies. In merged images, the parent-RFP and parent-GFP are false-colored in magenta and green, respectively. These controls are from the mutualism screen at 48 h (and are the controls shown in Fig. 5E). F) Five non-essential gene knockdowns exhibited mutualism. Top: competitive fitness of co-cultures with the wild-type-like parent-RFP (magenta) and the matrix production-deficient parent (red) at 16, 24, and 48 h. Numbers in top right indicate the mutualism score. Bottom: merged images of biofilm-colony co-cultures on MSgg-xylose agar at 48 h. Scale bar: 5 mm.

